# MOSAIC: A Joint Modeling Methodology for Combined Circadian and Non-Circadian Analysis of Multi-Omics Data

**DOI:** 10.1101/2020.04.27.064147

**Authors:** Hannah De los Santos, Kristin P. Bennett, Jennifer M. Hurley

**Affiliations:** Department of Computer Science, Rensselaer Polytechnic Institute, Troy, NY 12180, U.S.A.; Institute for Data Exploration and Applications, Rensselaer Polytechnic Institute, Troy, NY 12180, U.S.A.; Department of Mathematical Sciences, Rensselaer Polytechnic Institute, Troy, NY 12180, U.S.A.; Department of Biological Sciences, Rensselaer Polytechnic Institute, Troy, NY 12180, U.S.A.; Center for Biotechnology and Interdisciplinary Sciences, Rensselaer Polytechnic Institute, Troy, NY 12180, U.S.A.

## Abstract

**Motivation:** Circadian rhythms are approximately 24 hour endogenous cycles that control many biological functions. To identify these rhythms, biological samples are taken over circadian time and analyzed using a single omics type, such as transcriptomics or proteomics. By comparing data from these single omics approaches, it has been shown that transcriptional rhythms are not necessarily conserved at the protein level, implying extensive circadian post-transcriptional regulation. However, as proteomics methods are known to be noisier than transcriptomic methods, this suggests that previously identified arrhythmic proteins with rhythmic transcripts could have been missed due to noise and may not be due to post-transcriptional regulation.

**Results:** To determine if one can use information from less-noisy transcriptomic data to inform rhythms in more-noisy proteomic data, and thus more accurately identify rhythms in the proteome, we have created the MOSAIC (Multi-Omics Selection with Amplitude Independent Criteria) application. MOSAIC combines model selection and joint modeling of multiple omics types to recover significant circadian and non-circadian trends. Using both synthetic data and proteomic data from *Neurospora crassa*, we showed that MOSAIC accurately recovers circadian rhythms at higher rates in not only the proteome but the transcriptome as well, outperforming existing methods for rhythm identification. In addition, by quantifying non-circadian trends in addition to circadian trends in data, our methodology allowed for the recognition of the diversity of circadian regulation as compared to non-circadian regulation.

**Availability:** MOSAIC’s full interface is available at https://github.com/delosh653/MOSAIC.

**Supplementary information:** Supplementary data are available.at *Bioinformatics* online.

## 1 Introduction

Circadian rhythms are 24-hour endogenous cycles, reinforced by external cues. They allow an organism to optimize the timing of their cellular physiology to regulate biological processes in anticipation of the earth’s day/night cycle, thereby conferring an evolutionary advantage (Dunlap, 1999). Many processes, such as metabolic regulation, immune function, and sleep, are under the regulation of the circadian clock (Decoursey *et al*., 1997; Klarsfeld and Rouyer, 1998; Lévi *et al*., 2010; Ouyang *et al*., 1998). Chronic disruption of these rhythms is strongly associated with an increased risk of disease development, including cancer, diabetes, and cardiovascular disease (Evans and Davidson, 2013). At the molecular level, circadian rhythms are generated by a highly conserved circadian “clock” comprising a transcription-translation negative feedback loop on a 24-hour cycle (Hurley *et al*., 2016; Partch *et al*., 2014). While widespread regulation stemming from clock components is known at the transcriptional and translational levels, the complexity of the cellular circadian regulatory network is only beginning to be understood, e.g. (Hurley *et al*., 2014; Robles *et al*., 2014; Wang *et al*., 2017; Mure *et al*., 2018).

To determine what elements of cellular physiology are under the regulation of the circadian clock, large-scale “omics” datasets are gathered in order to quantify RNA expression (transcriptomics) or protein levels (proteomics) over time. Within these time courses, circadian rhythms manifest as oscillatory expression patterns, rising and falling throughout the day. As such, previous methods to identify circadian rhythms have compared changes in transcript or protein levels to reference fixed amplitude cosine waves (Wu *et al*., 2016; Hughes *et al*., 2010; Hutchison *et al*., 2015). However, these methods could not account for the prevalence of rhythms whose amplitudes change over time. This led to the development of the Extended Circadian Harmonic Oscillator model (ECHO), which introduced an amplitude change (AC) coefficient to the standard cosine model in order to capture these changing rhythms (De los Santos *et al*., 2020, 2019).

Recently, large-scale proteomics studies have shown that, though the central dogma of biology implies that rhythmic RNA would lead to rhythmic protein, transcript levels are often poorly correlated with protein levels, e.g. (Schwanhäusser *et al*., 2011). Moreover, we know from circadian proteomics analyses, e.g. Hurley et al. (2018), that rhythmic RNA expression does not necessarily imply rhythmic protein expression and vice versa (Hurley *et al*., 2018), These data suggest that there may be extensive post-transcriptional regulation in the clock’s output and beyond. The exact mechanisms of this regulation are currently unknown, though translation and degradation are predicted to be involved (Hurley *et al*., 2018; Lück *et al*., 2014; Collins *et al*., 2020).

In addition to the predicted sources of post-transcriptional regulation, it is known that the experimental methods used to gather proteomic data are noisier than those used to gather transcriptomic data (Hurley *et al*., 2018; Crowell *et al*., 2018). Thus, it is possible that circadianly-timed proteins may be missed due to higher background experimental noise rather than a true lack of circadian oscillation in the protein. However, if one could gain information from the underlying model for each gene in the transcriptome, one may be able to use that to bolster our confidence in the protein rhythmicity, or lack thereof, of the proteome (Misra *et al*., 2019; Subramanian *et al*., 2020). In the past, multiomics data has been used for the creation of biological networks in circadian biology and previous multiple omics studies have identified rhythms in each omics types separately before comparison, rather than jointly modeling them (Hurley *et al*., 2018; Hughes *et al*., 2009; Rund *et al*., 2011; Patel *et al*., 2012). This is likely due to the fact that, while many methods exist to identify circadian rhythms in a single omics type (e.g. (De los Santos *et al*., 2020; Hughes *et al*., 2010; Wu *et al*., 2016; Hutchison *et al*., 2015)), none exist that leverage multiple omics types to identify rhythms that may be masked by technical noise.

Further, though guidelines outlined by the circadian community for analyzing genome-scale experiments call for statistically quantifying arrhythmicity, traditional analyses of circadian rhythms in omics datasets do not quantify the significance of arrhythmic trends (Hughes *et al*., 2017). There is no wide-spread application for the analysis of model types beyond oscillatory models, as previous studies that utilized arrhythmic models applied their research only to single omics datasets with specifically tailored models (Wang *et al*., 2018; Hor *et al*., 2019; Keily *et al*., 2013). This suggests that the effects of significant rhythms can be overstated, as circadian-regulated processes that may also be controlled by arrhythmic genes cannot be quantified.

To address both of the above-described problems, we introduce MOSAIC (Multi-Omics Selection with Amplitude Independent Criteria). MOSAIC extends and augments the ECHO model by performing model selection on circadian omics data, including both non-circadian (linear and exponential) and circadian (ECHO and ECHO with a linear trend) models. MOSAIC then bridges multiple omics types through joint modeling, taking advantage of the transcriptome’s lower noise to identify rhythms previously not recovered in the proteome. Using both synthetic data and proteomic data from *Neurospora crassa*, we show that MOSAIC’s joint modeling approach accurately recovers rhythms at higher rates in the proteome. MOSAIC’s model selection also highlights the difference and extent of circadian regulation in different omics types. By accounting for universal trends, MOSAIC provides a more advantageous algorithm to allow for the mining of more accurate omics technologies (RNA-seq) to inform whether missed oscillations in less accurate omics technologies (proteomics, phosphoproteomics, metabolomics, etc.) level are due to technical noise or post-transcriptional regulation.

## 2 Methods

We created MOSAIC (Multi-Omics Selection with Amplitude Independent Criteria), which utilizes less-noisy omics data to inform rhythms in more-noisy omics data through a 4-stage workflow (Fig. 1). In stage 1, we identified the most probable model for each gene in the omics type, in this case the transcriptome and proteome, though other types of data are possible. We then moved to stage 2, where we jointly modeled both the proteome and the transcriptome based on those previously selected models. In stage 3, we then tested these new joint models against their previous independent models for improvement, and then moved to stage 4, where we evaluated goodness of fit for dataset analysis.

**Fig. 1.**
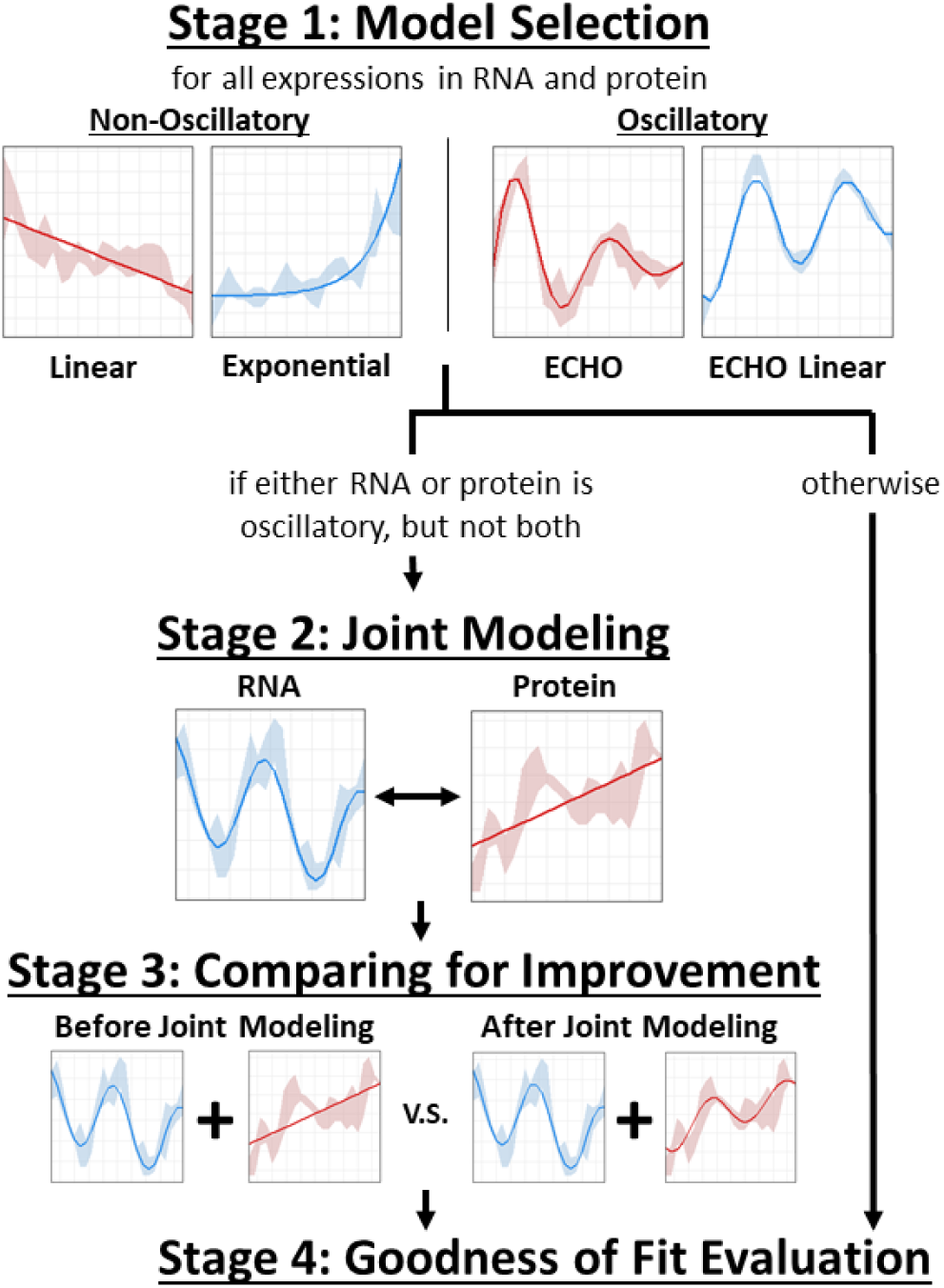
MOSAIC bridges multiple omics types to determine novel non-circadian and circadian trends. Overview of MOSAIC’s 4-stage workflow, comprised of Model Selection, Joint Modeling, Comparing for Improvement, and Goodness of Fit Evaluation. Data from (Hurley et al., 2014, 2018).

### 2.1 Stage 1: Model Selection

In our first stage, we identify the most probable models for each gene in each dataset. To do this, we select from a combination of oscillatory and non-oscillatory models:

Linear:

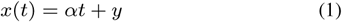

Exponential:

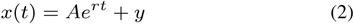

ECHO:

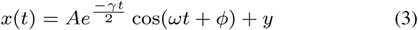

ECHO Linear:

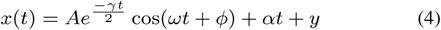

where the parameters are as follows: *x*(*t*) is the resulting change in output; *t* is time in hours; *α* is slope; *y* is equilibrium shift; *A* is initial amplitude; *r* is growth rate; *γ* is the amplitude change (AC) coefficient; *ω* is frequency; *ϕ* is phase shift. It should be noted that these models are nested, such that (1 - 3) are special cases of (4) with specific parameters set to 0.

Linear and exponential models are commonly used as examples of non-oscillatory data trends in the circadian literature (Deckard *et al*., 2013; De los Santos *et al*., 2020), while the ECHO model is a commonly used model for identifying circadian rhythms while taking into account amplitude change (De los Santos *et al*., 2020). We have extended the ECHO model to account for baseline changes over time, adding a linear term that was previously accounted for via preprocessing methods. Representative trends for each of these models appear in Supplemental Fig. 1.

By including both non-oscillatory and oscillatory models, we are able to encompass the majority of noted trends in circadian omics data (De los Santos *et al*., 2020; Wu *et al*., 2016; Deckard *et al*., 2013), providing for a more accurate comparison between both individual genes and the genes between different omics datasets. While constant models are absent from the model selection set, the prevalence of noise in biological data means that the slope of any omics data will realistically never be 0. Further, difficulty in estimating goodness of fit via p-value for constant models (as they are often the baseline implicit in null models) necessitates their disinclusion.

#### 2.1.1. Fitting Models to Experimental Data

To find the parameter values for (1, 2, 3, 4), we use the method of least squares. Given experimental data for each gene in each omics type **(t, x(t))** = (*t*_1_, *x*(*t*_1_)), (*t*_2_, *x*(*t*_2_)), …, (*t*_*n*_, *x*(*t*_*n*_)) and parameters *β*, the method of least squares minimizes the squared difference between experimental and fitted data as follows:

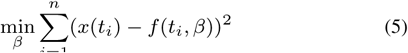

where *n* is the total number of data points and *f* (*t*_*i*_, *β*) refer to the equations and parameters in (1, 2, 3, 4). For the linear model, the parameters are *β* = (*α, y*); for the exponential, *β* = (*A, r, y*); for ECHO *β* = (*A, γ, ω, ϕ, y*); for ECHO Linear, *β* = (*A, γ, ω, ϕ, α, y*).

We use an ordinary linear least squares algorithm for the linear model (1), since it contains no nonlinear parameters. As this always results in the globally optimal solution, there is no need for the starting points required by the nonlinear method; we simply use the `lm` function in `R`.

All the other models, however, are nonlinear, necessitating the use of a nonlinear least squares algorithm. We find local solutions to this problem using the `nls.lm` algorithm, implemented `minpack.lm` package in `R`, which uses the Levenberg-Marquadt algorithm for nonlinear least squares.

Adjustments for multiple replicates for nonlinear equations are made as in (De los Santos *et al*., 2020), using weighted nonlinear least squares with weights at each time point equal to the inverse variance at each time point (Strutz, 2010). Due to the nonconvexity of our problem, it is important we choose the correct starting points. For each nonlinear model, we choose starting points and final fits for our parameters based on heuristics from the data and comparisons of different assumptions (Supplemental Section 2).

#### 2.1.2 Model Selection for Each Gene

After all models are fit, we choose the best model for each gene using the Bayesian Information Criterion (BIC) (Schwarz, 1978). The BIC for each fit is specified as the following for each model fit:

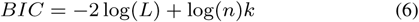

where *L* is the likelihood of the fit, *n* is the number of data points, and *k* is the number of parameters. We choose the model fit with the lowest BIC to represent the gene for the specific omics type. The two terms of the BIC rewards a higher likelihood and penalizes a more complex model respectively, providing a balance between choosing a highly parameterized model and overfitting.

### 2.2 Stage 2: Joint Modeling

Once the best model for each gene in each omics type is selected, we seek to address the noise in our multi-omics problem through joint modeling. We first determine whether joint modeling is appropriate for the current gene. If the gene is already oscillatory in both omics types, there is no need to use joint modeling to obtain new oscillatory parameters. If the gene is not oscillatory in either type, there is no basis for oscillations. Thus, we joint model genes that are oscillatory in only the transcriptome or the proteome.

If joint modeling is appropriate for a given gene, we use a joint model in which all parameters are allowed to be free for each omics type except for the period parameter. These are specified for each oscillatory model using the following equations:

ECHO Joint:

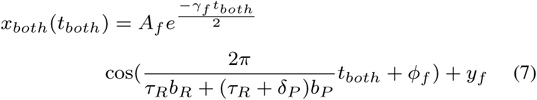

ECHO Linear Joint:

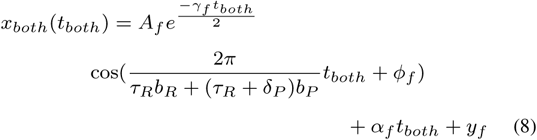

where *x*_*both*_(*t*_*both*_) is the concatenated experimental data for the transcriptome and proteome for a specified gene; *t*_*both*_ is the concatenated time points for the transcriptome and proteome; the subscript *f* indicates parameters which are independent for the transcriptome and proteome; *τ*_*R*_ and is the period, in hours, for the transcriptome; *δ*_*P*_ is the change in period, in hours, for the proteome; and *b*_*R*_ and *b*_*P*_ are either 1 or 0 depending on whether the time point corresponds to the transcriptome or the proteome, respectively. All other parameters retain their meaning from (3, 4).

The specification of this joint model largely stems from the fact that we want to “borrow” the oscillatory nature from one omics type and transfer it to the other. We force the proteome to maintain the same period, *τ*_*R*_, as the transcriptome, allowing for a slight deviation in the noisier proteome period. This deviation is kept within the resolution of the time points; i.e., if the time points had a resolution of 2 hours (hr), during fitting *δ*_*P*_ would be constrained to *±*2 hr. Through this joint modeling, We leave all other parameters independent between the two omics types, as we have no reason *a priori* why these should be kept the same.

We fit (7, 8) following the same procedure for nonlinear least squares (5) as described in Section 2.1.1. The final parameter fits of the joint model are obtained by comparing the results of two separate starting point regimes. The first results are obtained using the starting points derived from the averaged experimental data (Supplemental Section 2). The second results are obtained by using the results from a completely joint fitted model as starting points; that is, by eliminating *δ*_*P*_ in (7, 8). That completely joint model in turn uses the starting points to obtain its fit (Supplemental Section 2). In both starting point scenarios, *δ*_*P*_ is initially set to 0. The results of both starting point scenarios are chosen based on the fit with lowest AIC (Akaike, 1974). While leveraging these multiple starting points seems roundabout in theory, in practice, both starting points sets are chosen in relatively equal measure, yielding better fits overall.

### 2.3 Stage 3: Model Comparison

If joint modeling has been selected for the gene, we then move to determining whether the joint model produced a better parameter fit than either of the data fits individually. In order to determine this, we represent our individual models with parameters for the transcriptome and proteome fit separately as a joint model with completely free variables, where there is no dependence between the transcriptome and the proteome. This free representation allows for a correct comparison based on the amount of fitted data points. We then choose between these models using the BIC, selecting the joined model with the lowest BIC, as in Section 2.1.2.

### 2.4 Stage 4: Goodness of Fit Evaluation

Once we have chosen the best models to fit the experimental data, we need to determine whether the resulting fit approximates the data well. We estimate this goodness of fit by computing the p-value using Kendall’s tau rank correlation coefficient, which measures the concordance between two series of data (De los Santos *et al*., 2020; Hutchison *et al*., 2015). We use the p-value corresponding to the exact Kendall’s tau distribution. P-values calculated in this manner are reported for the transcriptomic and proteomic fits and the joint fit, if available. Further, if the selected model is linear, we also report the slope coefficient p-value, as calculated by the standard *t*-test by the `lm` function in `R`. All p-values are adjusted for multiple hypothesis testing using the Benjamini-Hochberg method and use this adjusted 0.05 cutoff, unless otherwise noted.

### 2.5 The MOSAIC Application

In order to facilitate ease of use, we have built the MOSAIC functionality into an app that allows users to find multi-omics trends and visualize the results, built using `R` Shiny (Wu *et al*., 2014) (Supplemental Section 3, Fig. 2). This interface is available on GitHub^1^. When running MOSAIC through this application, a variety of automatic preprocessing functionalities are available, including weighted smoothing, normalization, and removal of unexpressed genes (De los Santos *et al*., 2020) (Supplemental Fig. 2A). Further, specifications for period range (including the possibility of a “free run”, where no period range is specified) and changes to AC coefficient cutoffs for oscillatory models are also available. Automatic visualization for MOSAIC results include summary graphs for omics comparisons, gene expression plots, heat maps, and parameter density graphs (Supplemental Fig. 2B - F).

**Fig. 2.**
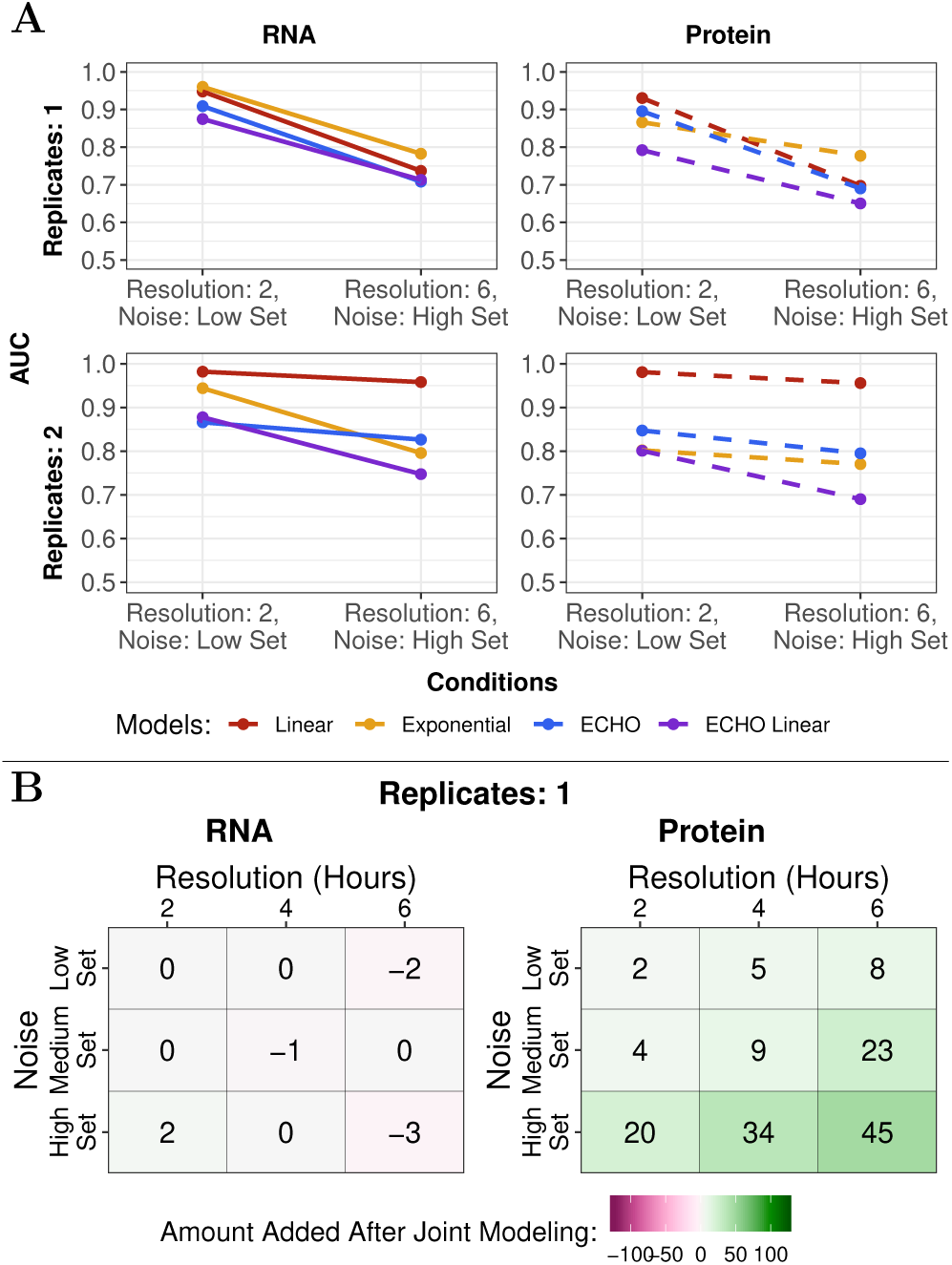
MOSAIC recovers models accurately regardless of data characteristics. A. AUCs for all models for synthetic transcriptome and proteome data at the best (2 hour resolution, low set noise) and worst (6 hour resolution, high set noise) sampling conditions for either one or two replicates. B. Heat map of how many oscillatory genes were recovered after the addition of joint modeling in MOSAIC, for varying noise and resolutions with 1 replicate using the synthetic transcriptomic and proteomic data. Annotations in each condition indicate the amount of oscillatory genes added after joint modeling.

## 3 Results

### 3.1 MOSAIC Accurately Recovers Disparate Models Through Model Selection and Joint Modeling

To evaluate MOSAIC’s effectiveness, we applied MOSAIC to generated synthetic datasets with each of our represented model types (Supplemental Section 4.1). We generated 12,000 genes for each omics type and condition, in a ratio of 2:1 non-circadian to circadian (8000 linear and exponential, 4000 of ECHO, ECHO linear, and their joint models) in order to mimic genome-wide datasets. These datasets were varied in sampling resolution (2, 4, 6 hours), replicates (1, 2, 3), and noise (low, medium, and high sets). To simulate the higher noise in proteomic data, we adjusted the synthetic “proteomic” data to have consistently higher noise than the “transcriptome” at all levels, hence creating noise “sets” (Supplemental Section 4.1.1).

To evaluate the efficacy of our modeling, we began by estimating AUCs for each model type using the ROC for the synthetic data (Fig. 2A, Supplemental Section 4.2, Tables 3 to 8). As expected, all models decreased in accuracy as noise increased and resolution/replication decreased, with larger decreases in the protein datasets. However, not all models decreased in accuracy at the same rate. Exponential models maintained the highest accuracy relative to other model classifications.

To explain this accuracy, we looked at the model misclassifications for each condition and omics type (Supplemental Section 4.2, Fig.s 4 to 6). At one replicate, the misclassification percentage increased with increasing noise and decreasing resolution at higher levels in the protein, as one would expect. However, several notable trends emerged. First, we noted that synthetic ECHO and ECHO Linear data were only misclassified into the other oscillatory model category, indicating that MOSAIC accurately identifies oscillatory models and explaining the supposedly low accuracy rate of ECHO linear models. Further, among most model types with the exception of exponential models, if the data was classified by the correct model, the BH-adjusted p-value was almost always below the 0.05 cutoff. With increasing replicates, these notable trends largely held and overall misclassification decreased.

We also investigated the ability of MOSAIC to jointly model and recover proteins by observing how many oscillating synthetic proteins were recovered before and after joint modeling at varying conditions (Fig. 2B, Supplemental Section 4.2, Fig. 7). With only one replicate, more oscillating proteins were recovered as noise increased and resolution decreased, with as many as 45 oscillating proteins rescued in the synthetic datasets, demonstrating the value of MOSAIC in high noise and low replicate situations (i.e. those found commonly in real omics data). These trends largely held with increasing replicates (Supplemental Fig, 7). However, when holding fixed resolution and varying the amount of replicates, we noted a boomerang effect; the amount of recovered proteins by joint modeling increased overall from one to two replicates, then decreased from two to three replicates. This phenomena suggests just enough information was added by two replicates to enhance joint modeling, but at three replicates, enough information was added to have high recovery in initial modeling. This explanation is bolstered by the overall counts of oscillatory genes before and after joint modeling in all scenarios. Surprisingly, though largely unaffected, at high noise levels, joint modeling was also able to rescue several oscillating transcripts. Overall, this demonstrated that joint modeling recovers large amounts of genes in the proteome, while losing little in the transcriptome.

### 3.2 MOSAIC Exceeds Common Methodologies in Recovering Circadian Rhythms

MOSAIC can be reduced to a circadian rhythm identification method by classifying its models as oscillatory and non-oscillatory. With this reduction, we used our synthetic data to compare MOSAIC to other commonly used methods for circadian rhythm identification: ECHO (De los Santos *et al*., 2020), JTK_CYCLE (JTK) (Hughes *et al*., 2010), and MetaCycle (Wu *et al*., 2016).

We first determined the F1-scores, a measure of accuracy ranging from 0 to 1, of all the methods at several BH-adjusted p-value cutoffs (Fig. 3A, Supplemental Section 4.2, Fig.s 8 to 10). The F1-scores of all methods decreased with increasing noise and resolution, though in different patterns. MOSAIC demonstrated high F1-scores regardless of condition, with a slight decrease with increasing sample resolution, though these decreases were mitigated by increasing replication. ECHO also maintained high F1-scores. JTK’s F1-scores were strongly impacted by increasing noise, and decreasing resolution and replication, decreasing to 0 as noise increased and resolution decreased to one replicate. Though not as strongly as JTK’s, MetaCycle’s F1-scores decreased in a similar manner.

**Fig. 3.**
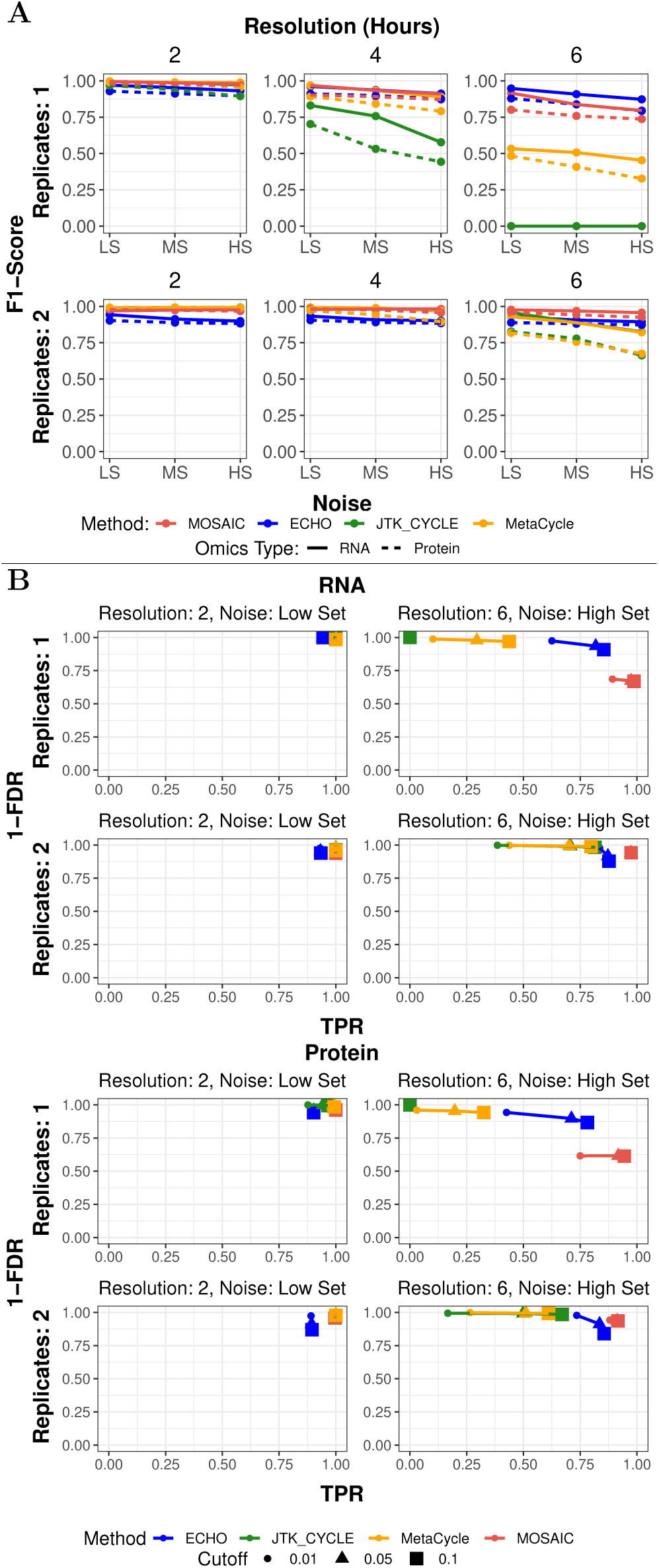
MOSAIC recovers more circadian genes than other analysis methods. A. F1-scores from several noise, resolution, and replicate levels in synthetic data for all tested methods, evaluated at a BH *p* of 0.05. B. Scatter plots of (1-FDR) versus TPR at three BH *p* cutoffs for synthetic transcriptome and proteome data, calculated at the best (2 hour resolution, LS) and worst (6 hour resolution, HS) sampling conditions for one and two replicates. LS = Low Noise Set, MS = Medium Noise Set, HS = High Noise Set.

To understand what accuracy tradeoffs we are making by using MOSAIC, we observed how BH-adjusted p-value cutoffs affected true positive rates (TPR) and false discovery rates (FDR) for each model (Fig. 3B, Supplemental Section 4.2, Fig.s 11 to 13). These rates were measured at different levels of noise, resolution, and replication. In general, all methods follow the pattern of decreasing overall accuracy as a balance of ECHO linear models showed the lowest accuracy rate. With increased replicates, accuracy increased throughout all models, though this was especially seen in linear model accuracy rates.

TPR and FDR as noise increases and resolution and replication decrease. Regardless of omics type, noise level, resolution, or replication, MOSAIC maintained a very high, if not perfect, TPR (Fig. 3B). However, increasing resolution had a strong impact on the MOSAIC FDR, though this effect was largely mitigated by increasing replicates. MOSAIC did not experience significant variation in TPR or FDR with increasing BH-adjusted p-value cutoffs, indicating low and stable p-values. While it did not have as high a TPR as MOSAIC, ECHO had a high TPR while maintaining a low FDR throughout all conditions (Fig. 3B). Increases in the BH-adjusted p-value cutoff were most impactful at high noise and low resolution/replication. JTK retained an extremely low FDR in all conditions (Fig. 3B). However, this came at the cost of a very low TPR, which decreased steeply with decreasing resolution and replication until the TPR became 0. MetaCycle largely followed JTK’s trends, though with a higher TPR (Fig. 3B). This shows that, at all conditions, MOSAIC recovers more true circadian genes than other methods in both omics types, with false positives in the worst conditions mitigated by increasing by even one replicate. As such, MOSAIC is the optimal method for the recovery of true circadian genes, with the caveat that this comes with the tradeoff of an increased FDR.

### 3.3 MOSAIC Increases Recovery and Understanding of Circadian Regulation in *Neurospora crassa*

To demonstrate MOSAIC’s efficacy on real data, we applied our method to publicly available transcriptomic and proteomic data from *Neurospora crassa* (Hurley *et al*., 2014, 2018) (Supplemental Section 4.3). Of the 4741 genes in common between the transcriptomic and proteomic datasets, prior to joint modeling MOSAIC found 4281 (90.3%) significant trends in the transcriptome and 2946 (62.1%) significant trends in the proteome. After joint modeling, MOSAIC found 4302 (90.7%) significant trends in the transcriptome and 3124 (65.9%) significant trends in the proteome, resulting in a total increase of 21 (0.4%) and 178 (3.8%) significant trends in the transcriptome and proteome respectively.

We next focused on joint modeling’s effects on increasing the identification of significant oscillatory trends. Before joint modeling, MOSAIC identified 3281 (69.2%) oscillating transcripts and 1132 (23.9%) oscillating proteins. After joint modeling, both omics types saw an increase in oscillatory trends, with joint modeling identifying 3324 (70.1%) oscillating transcripts and 1465 (30.9%) oscillating proteins, resulting in a total increase of 43 (0.9%) and 333 (7.0%) significant oscillatory trends in the transcriptome and proteome, respectively. Notably, we found increased identification not only at the proteomic level, but also in the transcriptome, suggesting that the incorporation of multiple omics types helps the recovery of oscillations from all types of omics data. Further, as there were more oscillatory rhythmic trends rescued than the total increase in significant trends, this suggests that the identification of oscillatory trends is bolstered by joint modeling.

When examining the overlap of circadian transcripts and proteins, we found that 1193 were oscillatory at both the transcriptomic and proteomic levels, while 2131 were oscillatory only in the transcriptome, and 272 were oscillatory only in the proteome, meaning that 18.5% of the identified proteome oscillates independently of the transcriptome. Hurley et al. (2018) found that 40% of the potential proteome oscillated independently of the transcriptome. Our joint modeling suggests that some of this discrepency came from noise in the proteomic data set. However, the maintenance of a 18.5% discrepancy between the transcriptome and proteome, despite recovering more genes than in the original proteomics study, solidifies the vital role of post-transcriptional regulation on circadian rhythms (Hurley *et al*., 2018).

We also noted that there were distinct distributions of model types between the transcriptome and proteome (Table 1). Of the 4302 transcripts whose trends we successfully modeled, the vast majority of the models were oscillatory (77.2%). By contrast, the proteins we successfully modeled had a much smaller percentage of oscillatory models (46.8%). Instead, the proteome was best modeled by linear and exponential models, with linear models representing the majority of non-oscillatory models. This discrepancy between the transcriptome and the proteome suggests that the circadian clock may more broadly regulate RNA as compared to protein.

**Table 1.** Model distributions suggest differential circadian regulation throughout the central dogma. Counts of significant model fits in transcriptomic and proteomic data from Neurospora crassa, as determined by MOSAIC, shows significant differences in the numbers of oscillating and non-oscillating transcripts and proteins.

|  | RNA (4302 total) | Protein (3124 total) |
| --- | --- | --- |
| Linear | 640 | 1148 |
| Exponential | 338 | 511 |
| ECHO | 1051 | 625 |
| ECHO Joint | 133 | 125 |
| ECHO Linear | 1883 | 465 |
| ECHO Linear Joint | 257 | 250 |

To further understand the difference in circadian regulation between the transcriptome and proteome, we performed gene ontological (GO) analysis (Supplemental Section 4.3). GO analysis revealed several notable trends in the distributions of biological process parent categories between non-circadian and circadian transcripts and proteins (Fig. 4). Principally, though metabolic processes and localization were the primary parent processes in all categories, oscillatory transcripts and proteins controlled a more diverse range of biological processes than non-oscillatory transcripts and proteins (Fig. 4A, B). Non-oscillatory transcripts were exclusively enriched in the parent processes of reproduction and developmental process, while oscillatory transcripts were more varied, with enrichment in parent processes related to response to stimulus, growth, rhythmic processes, and several others (Fig. 4A). In the proteome, parent processes exclusive to oscillating proteins showed a diverse set of physiological processes, including response to stimulus, rhythmic processes, reproduction, and multicellular organismal processes (Fig. 4B). Meanwhile, non-oscillating proteins were more limited in unique GO term categories, with only localization uniquely enriched.

**Fig. 4.**
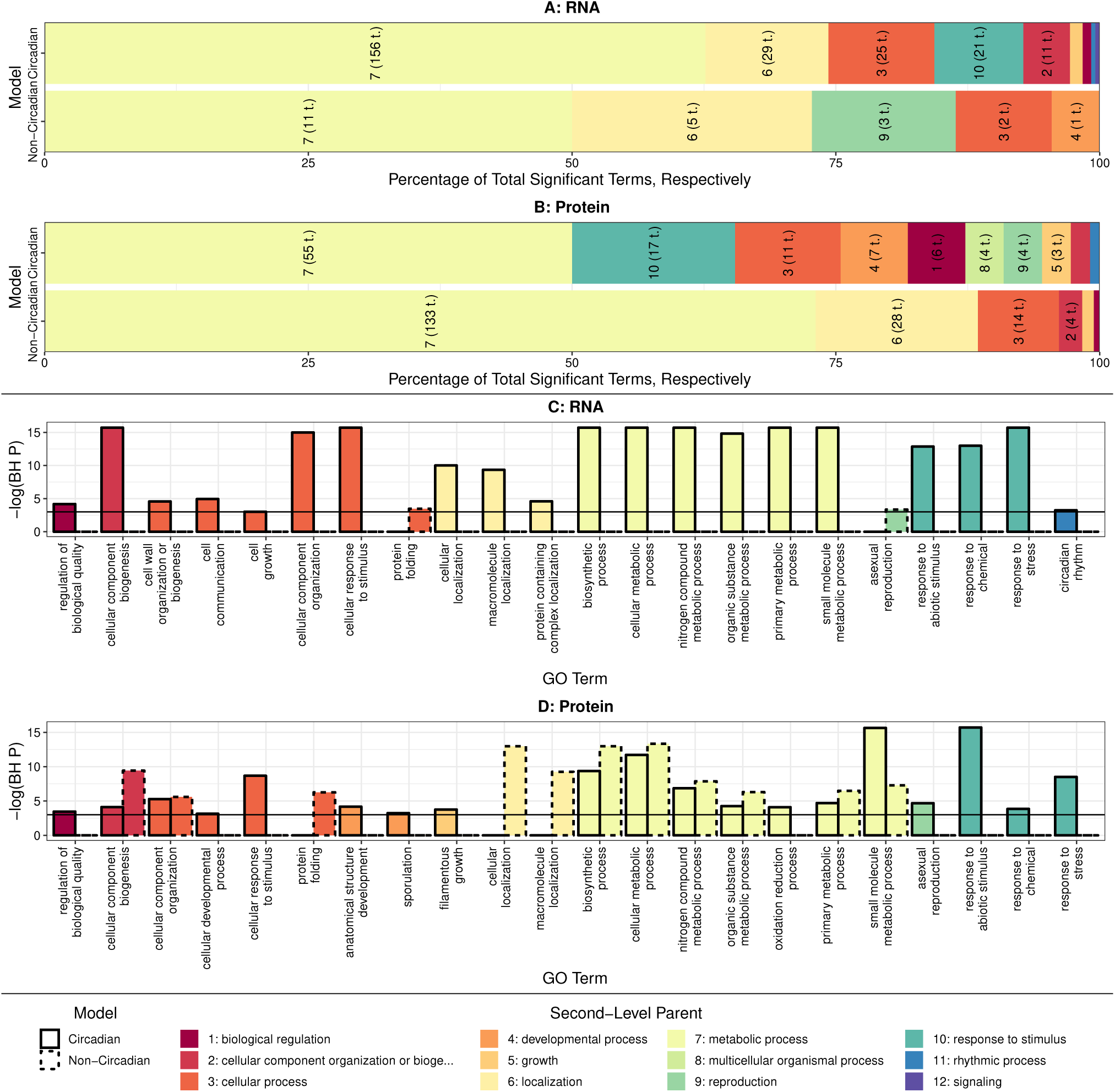
MOSAIC reveals distinct effects of circadian regulation between the Neurospora transcriptome and proteome. A and B. Percentages of significant GO terms, categorized by their second-level parent, for the circadian and non-circadian transcriptome (A) and proteome (B) in Neurospora crassa. Labeled numbers correspond to the second-level parent category, and numbers in parentheses indicate the total amount of significant terms corresponding to the category. C and D. Bar plots of negative log BH-adjusted p-values of third-level GO terms in transcriptome (C) and proteome (D), colored by second-level parent and with borders indicating circadian or non-circadian status.

Parental process distribution differences between non-oscillatory and oscillatory transcripts and proteins were further emphasized when examining the third-level children of the parent process (Fig. 4C and D). Oscillatory transcripts were exclusively enriched in many third-level children, including circadian rhythm (BH *p* = 3.9*e* − 2), response to stress (BH *p* = 9.7*e* − 9), and cellular component biogenesis (BH *p* = 2.1*e*−15) (Fig. 4C). Non-oscillatory transcripts were exclusively enriched only for asexual reproduction (BH *p* = 3.5*e* − 2). In the proteome, response to stress (BH *p* = 2.0*e* − 4), regulation of biological quality (BH *p* = 3.1*e* − 2), and asexual reproduction (BH *p* = 9.21*e* − 3) were uniquely enriched in oscillatory proteins (Fig. 4D). Non-oscillatory proteins were exclusively enriched in cellular localization (BH *p* = 2.3*e* − 6) and macromolecule localization (BH *p* = 9.6*e* − 5).

## 4 Discussion and Conclusion

We present MOSAIC, a novel method that can utilize joint modeling in multi-omics datasets to enhance the discovery of oscillating trends in circadian data, overcoming the noisiness that can hinder the discovery of oscillatory proteins in proteomic datasets (Fig. 1). Further, MOSAIC allows for the identification of non-oscillatory trends in omics data, thus providing a more-complete view of cellular regulation across circadian time in omics datasets. By building MOSAIC into an easy to use application, we have made this method accessible for any biologists interested in exploring the trends in their omics data over time (Section 3, Supplemental Fig. 2).

Synthetic data demonstrated MOSAIC’s efficacy in both model recovery and joint modeling (Fig. 2), as MOSAIC robustly recovered all potential models in the data and oscillatory gene recovery was enhanced by joint modeling (Fig. 2A, Supplemental Fig.s 4 to 6) 2B). Synthetic data also confirmed MOSAIC’s superior oscillatory trend identification, as MOSAIC retained higher F1-scores and true positive rates than other oscillatory identification methods (Fig. 3) (De los Santos *et al*., 2020;

Hughes *et al*., 2010; Wu *et al*., 2016). However, while false discovery rates remained comparable to other oscillatory identification methods for most conditions, at 6 hour resolution with one replicate, MOSAIC’s FDR significantly increased. We therefore recommend following the guidelines laid out in Hughes et al. (2017) as closely as possible when using MOSAIC (Hughes *et al*., 2017).

In real biological data, the analysis of transcriptomic and proteomic data from *Neurospora crassa* by MOSAIC identified over 300 novel oscillatory proteins as well as 43 oscillatory transcripts. This suggests that, despite higher overall noise levels in the proteome, jointly modeling the transcriptome and proteome can have a mutually beneficial relationship. This may be due to the fact that, while noise is generally higher in the proteome, it is not always higher in a one-to-one association, meaning a low noise protein could help a high noise transcript. (Supplemental Fig. 3). In addition, the nonconvexity of nonlinear least squares means that the formulation of a joint model may have allowed for the discovery of a lower local minimum that was not found when each of the omics types were specified separately.

MOSAIC’s use of non-oscillatory models also allows for the exploration of truly non-oscillatory elements in a dataset, not allowed for by other methods (De los Santos *et al*., 2020; Hughes *et al*., 2010; Wu *et al*., 2016). This permitted non-oscillatory enrichment beyond the absence of oscillatory enrichment. MOSAIC’s identification of both non-oscillatory and oscillatory trends illuminated a broader regulation of the trancriptome by the circadian clock as compared to the proteome in *Neurospora* (Table 1). Gene ontological analysis showed differences extended to biological output, where oscillatory genes regulated a more diverse set of processes than non-oscillatory genes (Fig. 4). While the difference in enrichment between omics types had been previously reported (De los Santos *et al*., 2019), the extension to non-circadian trends confirms this significant difference. Further, the use of MOSAIC’s joint modeling has confirmed that, while the extent is somewhat less than predicted, a discrepancy between oscillatory transcripts and proteins still existed, demonstrating the importance of post-transcriptonal regulation (Hurley *et al*., 2018).

In summary, we have shown MOSAIC to be a functional tool to identify oscillations masked by technical noise in multi-omics datasets through joint modeling. The ability of MOSAIC to identify both non-oscillatory and oscillatory trends allows for a fuller comprehension of circadian regulation. Though we here apply MOSAIC only to the transcriptome and proteome and investigate circadian biology, this multi-omics workflow could be easily extended to other omics types, e.g. phosphoproteome or metabolome, as well as other oscillatory processes, e.g. the cell cycle. As multi-omics circadian data becomes more prevalent (Collins *et al*., 2020; Campbell *et al*., 2020; Hughes *et al*., 2017), MOSAIC will provide an important role in finding and understanding both oscillatory and non-oscillatory trends in a variety of organisms and processes.

## Acknowledgements

We would also like to thank Emily Collins, Gretchen Clark, and Meaghan Jankowski for their biological expertise, as well as their advice regarding the MOSAIC application.

## Footnotes

1 https://github.com/delosh653/MOSAIC

## References

Akaike, H. (1974). A new look at the statistical model identification. IEEE Trans. Autom. Control, 19(6), 716–723.

Campbell, K. et al. (2020). Building blocks are synthesized on demand during the yeast cell cycle. Proceedings of the National Academy of Sciences, page 201919535.

Collins, E.J. et al. (2020). Post-transcriptional circadian regulation in macrophages organizes temporally distinct immunometabolic states. bioRxiv.

Crowell, A.M. et al. (2018). Learning and imputation for mass-spec bias reduction (LIMBR). Bioinf., 35(9), 1518–1526.

De los Santos, H. et al (2019). ENCORE: A visualization tool for insight into circadian omics. In Proc. of the 10th ACM Int. Conf. on Bioinf., Comput. Biol. and Health Inform., ACM-BCB ‘19, New York, NY, USA. ACM.

De los Santos, H. et al (2020). ECHO: an application for detection and analysis of oscillators identifies metabolic regulation on genome-wide circadian output. Bioinf., 36(3), 773–781.

Deckard, A. et al. (2013). Design and analysis of large-scale biological rhythm studies: a comparison of algorithms for detecting periodic signals in biological data. Bioinf., 29(24), 3174–3180.

Decoursey, P.J. et al. (1997). Circadian performance of suprachiasmatic nuclei (scn)-lesioned antelope ground squirrels in a desert enclosure. Physiol. Behav., 62(5), 1099—-1108.

Dunlap, J.C. (1999). Molecular bases for circadian clocks. Cell, 96(2), 271–290.

Evans, J.A. and Davidson, A.J. (2013). Health consequences of circadian disruption in humans and animal models. In Progress in Molecular Biology and Translational Science, volume 119, pages 283–323. Elsevier.

Hor, C.N. et al. (2019). Sleep–wake-driven and circadian contributions to daily rhythms in gene expression and chromatin accessibility in the murine cortex. Proc. Natl. Acad. Sci., 116(51), 25773–25783.

Hughes, M.E. et al. (2009). Harmonics of circadian gene transcription in mammals. PLoS Genet., 5(4), e1000442.

Hughes, M.E. et al. (2010). JTK_CYCLE: An efficient nonparametric algorithm for detecting rhythmic components in genome-scale data sets. J. Biol. Rhythms, 25(5), 372–380.

Hughes, M.E. et al. (2017). Guidelines for Genome-Scale Analysis of Biological Rhythms. J. Biol. Rhythms, 32(5), 380–393.

Hurley, J.M. et al. (2014). Analysis of clock-regulated genes in Neurospora reveals widespread posttranscriptional control of metabolic potential. Proc. Natl. Acad. Sci., 111(48), 16995–17002.

Hurley, J.M. et al. (2016). Circadian oscillators: Around the transcription–translation feedback loop and on to output. Trends Biochem. Sci., 41(10), 834–846.

Hurley, J.M. et al. (2018). Circadian proteomic analysis uncovers mechanisms of post-transcriptional regulation in metabolic pathways. Cell Syst., 7(6), 613–626.e5.

Hutchison, A.L. et al. (2015). Improved statistical methods enable greater sensitivity in rhythm detection for genome-wide data. PLoS Comput. Biol., 11(3), e1004094.

Keily, J. et al. (2013). Model selection reveals control of cold signalling by evening-phased components of the plant circadian clock. Plant J., 76, 247–257.

Klarsfeld, A. and Rouyer, F. (1998). Effects of circadian mutations and ld periodicity on the life span of drosophila melanogaster. J. Biol. Rhythms, 13(6), 471—-478.

Lévi, F. et al. (2010). Circadian timing in cancer treatments. Annu. Rev. Pharmacol. Toxicol., 50(1), 377–421.

Lück, S. et al. (2014). Rhythmic degradation explains and unifies circadian transcriptome and proteome data. Cell Reports, 9(2), 741–751.

Misra, B.B. et al. (2019). Integrated omics: tools, advances and future approaches. Journal of Molecular Endocrinology, pages R21–R45.

Mure, L.S. et al. (2018). Diurnal transcriptome atlas of a primate across major neural and peripheral tissues. Science, 359(6381), eaao0318.

Ouyang, Y. et al. (1998). Resonating circadian clocks enhance fitness in cyanobacteria. Proc. Natl. Acad. Sci., 95(15), 8660–8664.

Partch, C.L. et al. (2014). Molecular architecture of the mammalian circadian clock. Trends Cell Biol., 24(2), 90–99.

Patel, V.R. et al. (2012). CircadiOmics: integrating circadian genomics, transcriptomics, proteomics and metabolomics. Nat. Methods, 9(8), 772–773.

Robles, M.S. et al. (2014). In-vivo quantitative proteomics reveals a key contribution of post-transcriptional mechanisms to the circadian regulation of liver metabolism. PLoS Genet., 10(1), e1004047.

Rund, S.S.C. et al. (2011). Genome-wide profiling of diel and circadian gene expression in the malaria vectorAnopheles gambiae. Proc. Natl. Acad. Sci., 108(32), E421–E430.

Schwanhäusser, B. et al. (2011). Global quantification of mammalian gene expression control. Nature, 473(7347), 337–342.

Schwarz, G. (1978). Estimating the dimension of a model. Ann. Stat., 6(2), 461–464.

Strutz, T. (2010). Data Fitting and Uncertainty: A Practical Introduction to Weighted Least Squares and Beyond. Vieweg and Teubner, Germany.

Subramanian, I. et al. (2020). Multi-omics data integration, interpretation, and its application. Bioinformatics and Biology Insights, 14, 117793221989905.

Wang, J. et al. (2017). Nuclear proteomics uncovers diurnal regulatory landscapes in mouse liver. Cell Metab., 25(1), 102–117.

Wang, J. et al. (2018). Circadian clock-dependent and -independent posttranscriptional regulation underlies temporal mRNA accumulation in mouse liver. Proc. Natl. Acad. Sci., 115(8), E1916–E1925.

Wu, G. et al. (2014). Evaluation of five methods for genome-wide circadian gene identification. J. Biol. Rhythms, 29(4), 231—-242.

Wu, G. et al. (2016). MetaCycle: an integrated r package to evaluate periodicity in large scale data. Bioinf., 32(21), 3351–3353.

