## Supplemental Document for "MOSAIC: A Joint Modeling Methodology for Combined Circadian and Non-Circadian Analysis of Multi-Omics Data"

April 23, 2020

[1]Department of Computer Science, Rensselaer Polytechnic Institute, Troy, NY, U.S.A
[2]Institute of Data Exploration and Applications, Rensselaer Polytechnic Institute, Troy, NY, U.S.A
[3]Department of Mathematical Sciences, Rensselaer Polytechnic Institute, Troy, NY, U.S.A
[4]Department of Biological Sciences, Rensselaer Polytechnic Institute, Troy, NY, U.S.A
[5]Center for Biotechnology and Interdisciplinary Sciences, Rensselaer Polytechnic Institute, Troy, NY, U.S.A

#### 1 Representative Trends for Each Model Type

Representative trends for each model type appear in Figure 1.

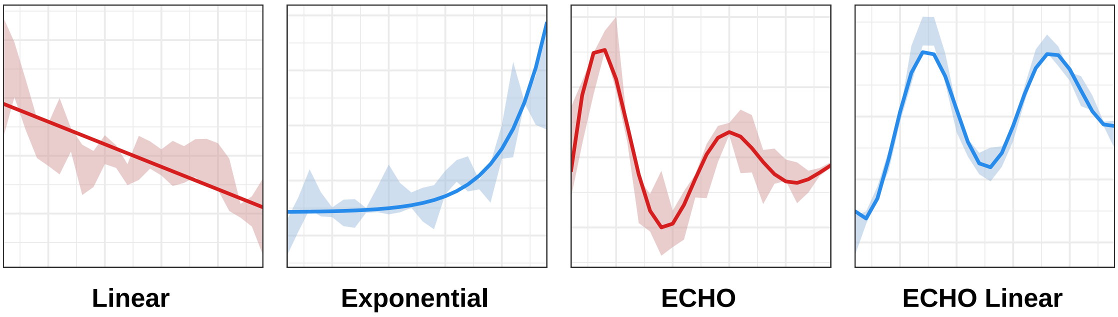


Figure 1: **MOSAIC includes both oscillatory and non-oscillatory models.** Example curves for each model MOSAIC considers during model selection. Non-oscillatory models are linear and exponential models. Oscillatory models are ECHO and ECHO Linear models. Red curves indicate proteomic data, and blue curves indicate transcriptomic data. Data from Hurley et al. (2014) and Hurley et al. (2018) [9, 10].

#### 2 Starting Points and Fitted Parameters for Nonlinear Models

To find the starting points for the nonlinear least squares problem for our nonlinear models (M.1 - M.4)[1] (Section M.2.1.1), we developed novel initialization schemes and enhanced older methods. Starting points (denoted with subscript 0), regardless of model, were based on the average of any replicates in the experimental data (denoted by $\overline{\mathbf{x(t)}}$).

[1]Elements from the main text are referred to by M.X, where X is the element number.

#### 2.1 Exponential Model Procedures

For the exponential model (M.2), starting points were obtained based on the idea that the log of an exponential model (with no equilibrium shift) produces a straight line, as shown by the following:

$$\begin{aligned} \log(Ae^{rt}) &= \log(A) + \log(e^{rt}) \\ &= \log(A) + rt \end{aligned} \tag{1}$$

where all parameters maintain their notation from (M.2). Thus, with exponential data, it is possible to get the initial amplitude and growth rate by performing linear least squares on the log of the data. However, two points preclude this simplicity when applied to our experimental data. First, our model has an equilibrium shift, which we need to remove before estimating our parameters with linear least squares. Second, log restricts the estimated initial amplitude to remain positive, which is not necessarily true in real data. As such, we need to pursue two assumptions: positive initial amplitude, where our data remains as estimated; and negative initial amplitude, where our data is negated. We take these points into account when estimating our initial starting points.

Our negative and positive initial amplitude schemes require slightly different initial transformations to the averaged data as follows:

$$\mathbf{x}_{tf,pos} = \overline{\mathbf{x(t)}} - \min_t(\overline{\mathbf{x(t)}}) \tag{2}$$

$$\mathbf{x}_{tf,neg} = -\overline{\mathbf{x(t)}} - \min_t(-\overline{\mathbf{x(t)}}) \tag{3}$$

where $\mathbf{x}_{tf}$ is the transformed averaged data, the *pos* subscript indicates results related to the positive initial amplitude assumption, and the *neg* subscript indicates the negative.

For both assumptions, the resultant 0 of $\mathbf{x}_{tf}$ (corresponding to the minimum of $\pm\overline{\mathbf{x(t)}}$, respectively) was then replaced by the minimum of the nonzero entries of $\mathbf{x}_{tf}$. This replacement allowed us to not skew the log of these points with an outlier, as $\log(0) = -\inf$.

Then, to find the initial amplitude ($A_0$) and growth rate ($r_0$), we performed the following ordinary least squares problem:

$$\min_{A_{\log},r} \sum_{i=1}^{n} (\log \overline{x}_{tf}(t_i) - (r_f t_i + A_{\log}))^2 \tag{4}$$

where $A_{\log}$ is the log of the fitted initial amplitude, $r_f$ is the fitted growth rate, $n$ is the total number of data points in the averaged data, and $x_{tf}(t)$ is the transformed data. Thus, by this method, the starting points for the exponential model with both the positive and negative assumptions are:

$$A_{0,pos} = \exp(A_{f,log,pos})$$

$$r_{0,pos} = r_{f,pos}$$

$$y_{0,pos} = \min(\overline{\mathbf{x(t)}})$$

$$A_{0,neg} = -\exp(A_{f,log,neg})$$

$$r_{0,neg} = r_{f,neg}$$

$$y_{0,pos} = -\min(-\overline{\mathbf{x(t)}})$$

After obtaining these starting points, we fit (M.2) using nonlinear least squares (M.5) for the positive and negative initial amplitude assumptions separately. Thus, we obtain two final parameter fits corresponding to each assumptions. We then choose between these parameter fits with the AIC, choosing the fit yielding the lowest AIC [1].

#### 2.2 ECHO Model Procedures

For the ECHO model (M.3), starting points are similar to those specified in [6], with some enhancements to both the starting points and the data. First, the averaged data is smoothed with a 1-2-1 weighting scheme described in [6]. Then starting points are calculated based on the smoothed averaged data as follows:

$$A_0 = \begin{cases} |\text{peaks}(1)|, & \text{if } \# \text{ of peaks} > 0 \\ \max(\overline{x_s(t)}) - y_0, & \text{otherwise} \end{cases} \tag{5}$$

$$\gamma_0 = \begin{cases} \frac{1}{\sqrt{1+(\frac{2\pi}{\delta})^2}}, & \text{if } \# \text{ of peaks } \geq 2 \ \& \ \text{peak}(1) \ > \ \text{peak } (2) \\ \frac{-1}{\sqrt{1+(\frac{2\pi}{\delta})^2}}, & \text{if } \# \text{ of peaks } \geq 2 \ \& \ \text{peak}(1) \ < \ \text{peak } (2) \\ 0.01, & \text{if peaks } = 1 \\ 0, & \text{otherwise} \end{cases} \tag{6}$$

$$\omega_0 = 2\pi / \begin{cases} (\# \text{ of time points})(\text{resolution})(\# \text{ of peaks}), & \text{if } \# \text{ of peaks} > 1 \\ (\# \text{ of time points})(\text{resolution})(\# \text{ of peaks} + 1), & \text{if } \# \text{ of peaks} = 1 \\ (\# \text{ of time points})(\text{resolution})(H + L)/2, & \text{if } \# \text{ of peaks} = 0 \end{cases} \tag{7}$$

$$y_0 = \overline{\overline{x_s(t)}} \tag{8}$$

where # is number, $\overline{x_s(t)}$ is the smoothed averaged data, and resolution is the difference, in hours, between time points. Similarly to [6], we calculate $\phi_0$ by partitioning the possible phase range into 12 parts such that $\phi_0 = \frac{i\pi}{6}, i = 0, ..., 11$. Then, using the other previously calculated initial values, we choose the $\phi_0$ that minimizes the absolute value of the the difference between $\overline{x_s(t)}$ and proposed initial fit at certain time points. The compared values are calculated at time points from the beginning, middle, and end of the time course: time points 2 and 3, $\lfloor\frac{(\# \text{ of time points})}{2}\rfloor$ and $\lfloor\frac{(\# \text{ of time points})}{2}\rfloor + 1$, and $(\# \text{ of time points}) - 2$ and $(\# \text{ of time points}) - 1$.

Equations (5, 6, 7) depend on the peaks vector, which is calculated from $\overline{x_s(t)}$. A value is determined to be a "peak" by being either the maximum of minimum of a set consisting of the value and its surrounding points; i.e., by being a peak or a trough. These adjacent surrounding points are the same as those stated in [6].

The final vector of "peaks" is calculated by comparing the vector of all peaks and all troughs found across the time course for consistency, defined by the following:

$$C = -|\tau_{est} - \tau_{eff}| \tag{9}$$

$$\tau_{est} = \frac{\text{time course length(hours)}}{\#\text{of peaks}} \tag{10}$$

$$\tau_{eff} = \text{mean}_{t_j > t_i}(\text{peak}_j - \text{peak}_i) \tag{11}$$

where $C$ is consistency, $\tau_{est}$ is estimated period, $\tau_{eff}$ is effective period, and a "peak", in this notation, corresponds to a peak or a trough. Consistency is calculated for both the set of peaks and the set of troughs.

If troughs have a higher consistency than peaks, or there is one peak or less and the number of troughs is greater than the number of peaks, then the troughs are selected for the "peaks" vector. Otherwise, the peaks are selected. By changing this "peaks" vector calculation, we more accurately account for changes in phase and mitigate noise effects.

The calculation of the logarithmic decrement, as included in (6), has been amended to the following:

$$\delta = \begin{cases} \delta = \ln(\frac{x_s(t)}{x_s(t+T)}) & \text{if peak}(1) \ > \ \text{peak } (2) \\ \delta = \ln(\frac{x_s(t+nT)}{x_s(t)}) & \text{otherwise} \end{cases} \tag{12}$$

where $t$ is the time of the first peak and $t + T$ is the time of the next peak. The previous iteration of the logarithmic decrement underestimated the effects of damping or forcing, respectively.

After all initial parameters are calculated, ECHO's starting points are then fit using nonlinear least squares (M.5) as in [6], with one crucial difference. The AC coefficient, $\gamma$, is restricted to be

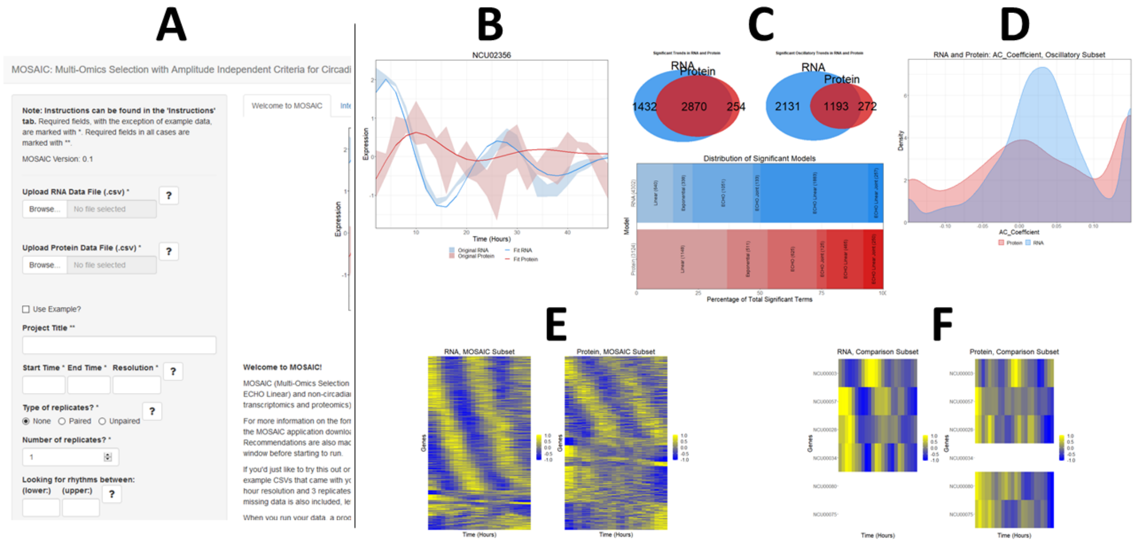


Figure 2: **The MOSAIC application provides an easy-to-use interface for circadian biologists to run the algorithm and visualize corresponding results.** Under the "Finding Trends" part of MOSAIC (A), users can upload data, specify data characteristics, and choose from various preprocessing methods. In the "Visualizing Results" part, users can generate Gene Expression Plots (with/without replicates) (B), Summary Visualizations (C), Parameter Density Graphs (D), Heat Maps (E), and Comparison Heat Maps (F), described in Section 3. Data visualized is transcriptomic and proteomic *Neurospora crassa* data, processed as described in 4.3 [9, 10].

within the range of damped, harmonic, and forced, excluding overexpression and repression. As overexpressed and repressed genes are not considered oscillatory, we excluded them from MOSAIC analysis, as non-oscillatory genes are accounted for by this method. Recommended $\gamma$ cutoffs for overexpressed and repressed genes are used in MOSAIC, unless specified otherwise.

##### 2.3 ECHO Linear Model Procedures

For the ECHO Linear model (M.4), starting points are calculated in the same manner as the ECHO model, with compensation for the linear term. Initial slope $\alpha_0$ and equilibrium shift $y_0$ are calculated with ordinary least squares on the total experimental data. The estimated linear trend is removed from the averaged data, and then all parameters are calculated and fit as in the ECHO model.

#### 3 MOSAIC Application Features

The MOSAIC Application[2] is divided into 2 parts: finding trends and visualizing results (Fig. 2). In the first part, Finding Trends, users upload their transcriptomic and proteomics datasets, specifying pertinent information about the dataset, including the time range (in hours), time point resolution, and number of replicates (Fig. 2A). The user can then select from three preprocessing methods: smoothing with a 1-2-1 weighting, z-score normalization, and removal of unexpressed genes. It is important to note that preprocessing affects parameter estimation. Users can also specify a range of periods (or a free run) and any changes to AC coefficient cutoffs for oscillatory models. These preprocessing methods and their effects, as well as free run specifications, are discussed in [6]. Results are downloadable as a CSV or as an .RData file. The .RData file contains both the results and metadata about the MOSAIC run, and is used for the next part of the application, Visualizing Results. By creating a simple interface by which to run MOSAIC, we make this powerful method more widely available to circadian biologists, more accessible than the traditional code script that pervades the field.

After uploading MOSAIC .RData results to Visualizing Results, users can choose from several different automatic visualizations: Summary Visualization, Gene Expression Plots, Heat Maps,

[2]https://github.com/delosh653/MOSAIC

Comparison Heat Maps, and Parameter Density Graphs (Fig. 2B - F). Throughout, color coding designates RNA as shades of blue and protein as shades of red. These visualizations, as a whole, make sifting through results more accessible and easier for users, such that they will be able to derive omics insights more quickly and easily.

In the Summary Visualization, users can compare resulting significant model distributions from transcriptomics (RNA) and proteomics (protein) (Fig. 2C). At the top of the visualization, the left Venn Diagram shows the overlap of significant models in RNA and protein, while the right shows the overlap of significant oscillatory models. Below this, a comparative bar graphs shows the percentages of each significant model relative to the total number of significant models in RNA or protein, respectively. Counts of significant models are also displayed on the bar graph alongside model and omics type names. These counts are also reflected below the visualization, along with additional categories such as AC coefficient categories. By introducing MOSAIC's output through a Summary visualization, users are more able to compare and contrast the differences between different omics types. These differences may stem from distributions of the amounts of models, as illustrated by the comparative bar graphs, or in the significant and oscillatory genes they hold in common, as illustrated by the Venn Diagrams. As such, the user gains a comprehensive view of MOSAIC results with ease, which further research objectives.

Gene Expression Plots display processed gene expression values and fitted models for a selected genes (Fig. 2B). The fitted model is displayed as the dark corresponding color, while original data is displayed either as a lighter shaded line or, if replicates exist, filled in between the maximum and minimum replicate value at each time point. The ability to plot replicates, if they exist, is available as well. Expressions for RNA and protein can be plotted overlaid with each other, side-by-side in separate plots, or plotted with only one or the other on the plot. Below the visualization, relevant model parameters and p-values are listed. These plots and parameters, particularly in their ability to be overlaid, allow users to dig down to the differences in transcriptomics and proteomics on the gene level. The comparisons of these models in two omics types allow users to understand post-transcriptional regulation on a gene scale.

Heat Maps visualize averaged expression for a selected subset of genes (Fig. 2E). The expression for each gene in the heat map is mean-centered and normalized by the absolute maximum value of the averaged replicate expression, such that each row is on a scale of $[-1, 1]$. Individual replicates can also be visualized. High expression values are colored yellow, while low values are colored blue. Available subsets for the heat map include the MOSAIC subset, each model subset, and AC coefficient subsets, among others. Heat maps are ordered first by model type, then by specified parameters, in the following manner: oscillatory genes, sorted by phase; exponential genes, subset by growth rate and then amplitude, ordered by growth rate; linear genes, ordered by slope. Heat maps for RNA and protein can be displayed side-by-side or individually. Total amounts of genes for each heat map are displayed below the plot. The inclusion of heat maps in the MOSAIC application allows users to see significant gene expression overall, as well as the prevalence significant model types for each omics type. This side-by-side expression comparison furthers the observation of differences between omics types.

Heat Map Comparison plots are an extension of heat maps, which provide a method for direct comparison of user supplied lists of genes (Fig. 2F). Users enter genes for comparison in both RNA and Protein, which are displayed in heat maps side-by-side, using white tiles in each heat map to reflect genes that were only specified in the opposite omics type subset. These heat maps are ordered by a user-specified dominant omics type. This applies heat map ordering described above to the dominant omics type, then uses this exact ordering on both omics types. Below the plot, gene names for those that appear in both and only each omics types are displayed. Much like the Heat Maps and Gene Expression Plots, these comparisons allow the viewer to more accurately compare gene expression between omics types. By placing these heat maps side by side, users can more accutely see the difference in average expression for the specified subset. These subsets can answer research questions, such as, for example, a subset relating to specific biological functions, and show how those functions manifest overall on a gene expression scale.

In Parameter Density Graphs, density graphs for a specified parameter and subset are displayed, showing the range and distribution of the specified parameter (Fig. 2D). RNA and protein graphs can be overlaid, plotted side-by-side, or plotted separately. Below the plot, quantile information for each distribution appears. By providing parameter density plots, users can compare the distributions of parameters for different omics types, viewing how the specified subgroup biases certain parameters, such the AC coefficient or slope, depending on omics type.

| Parameter Name | Parameter Symbol | Range |
|---|---|---|
| Equilibrium Shift | $y$ | $[-4, 4]$ |
| Slope | $\alpha$ | $\pm[0.05, 1]$ |
| Growth Rate | $r$ | $\pm[0.05, 0.25]$ |
| Initial Amplitude, Exponential | $A$ | $\pm[0.01, 5]$ |
| Initial Amplitude, Oscillatory | $A$ | $\pm[8, 10]$ |
| Amplitude Change (AC) Coefficient | $\gamma$ | $[-0.15, 0.15]$ |
| Radian Frequency | $\omega$ | $2\pi/[20, 28]$ |
| Phase Shift | $\phi$ | $[-2\pi, 2\pi]$ |

Table 1: Parameter ranges for synthetic data generation.

Additional auxiliary information is displayed in separate tabs, giving users further access to concise results and information. Gene lists display gene names and parameter information for specified subsets in RNA and Protein. This information is displayed side-by-side for both RNA and protein, overlaid, or for each individually. Overlaid gene lists indicate a subset where criteria is true for RNA or protein. User inputs for the MOSAIC run appear in another tab, including information such as data inputs and version information.

### 4 Data

#### 4.1 Synthetic Dataset Generation

In order to test MOSAIC's effectiveness, we generated several synthetic data sets to represent transcriptomic and proteomic data at various resolutions, noise levels, and replicates, enhancing the data simulation generations proposed in [18] and [7]. For each transcriptomic and proteomic dataset, we generated 12000 samples total. 4000 of these were of our non-oscillatory models (linear (M.1) and exponential (M.2)) and 1000 each were of our oscillatory models (ECHO (M.3), ECHO Joint (M.4), ECHO Linear (M.7), and ECHO Linear Joint (M.8)). This created an 8000:4000, or 2:1, nonoscillatory to oscillatory ratio, so as to simulate both the size and scale of a genome-wide dataset. Parameters for all models were randomly drawn from uniform distributions with ranges specified in Table 1, with the exception of $\delta_P$, which was drawn from a normal distribution with mean 0 and standard deviation $\frac{2}{3}$.

Time courses for each simulated data set were varied in resolution, noise levels, and replicates. Resolution was varied over 2, 4, and 6 hours over a 48-hour time course, created through subsampling. Replicates were varied at 1, 2, and 3 samples at each time point. Noise levels varied between low, medium, and high "sets", described in Section 4.1.1.

##### 4.1.1 Generating Noise for Synthetic RNA and Protein

Due to the differences in overall noise levels in RNA and protein and the ambiguity of noise additions to nonoscillatory models in previous studies, we have developed a new method of creating synthetic omics data in order to more accurately simulate real noise. Previously, noise was calibrated based on the initial amplitude of the fixed amplitude oscillators being tested [18, 7]. However, as we know from various studies of rhythms with changing amplitude [6, 17], initial amplitude is not always effective amplitude; for example, the initial amplitude of a forced rhythm increases over time. We also know the standard deviation of Gaussian noise applied in previous studies is often fixed, which does not reflect the noise effects we see in real data, as overall noise levels in expressions can even vary from gene to gene [12]. Further, non-oscillatory models such as linear models do not have initial amplitudes.

To address these problems, we have created a two-step method for generating noise more similar to real data, regardless of the underlying model. First, we redefine noise to be a percentage of amplitude rather than a fixed additive amount for all models. Noise for each gene and sample is drawn from a normal distribution centered at 0 and a standard deviation for each gene. The standard deviation distribution, from which the gene's standard deviation is drawn, is normally

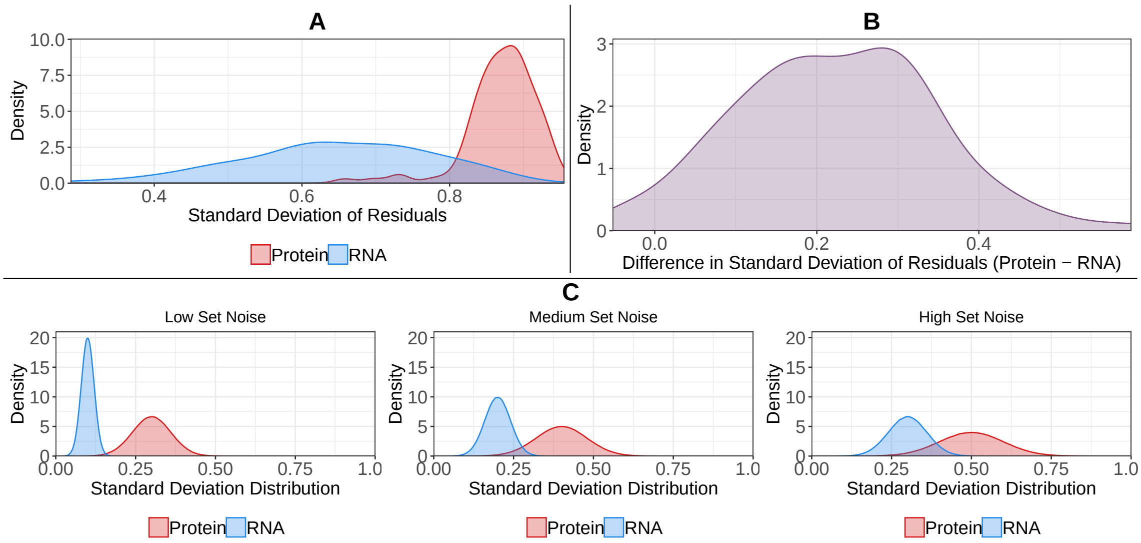


Figure 3: ***Neurospora crassa* standard deviation distributions inform synthetic noise distributions [9, 10].** A. Density plot of standard deviations of residuals for the transcriptome (RNA) and proteome (protein). These are residuals of significant oscillatory models, as determined by MOSAIC on normalized *Neurospora* data. B. Density plot of difference between standard deviations of protein and RNA, where a negative value indicates a higher RNA standard deviation and a positive value indicates a higher protein value. C. Example RNA and protein noise level distributions for an amplitude of 1 and mean standard deviation for the level (Table 2).

| | **Transcriptome** | **Proteome** |
|---|---|---|
| **Low Set** | 1/10 | 3/10 |
| **Medium Set** | 2/10 | 4/10 |
| **High Set** | 3/10 | 5/10 |

Table 2: **Noise levels in synthetic data reflect the noise difference in transcriptome and proteome.** Mean standard deviations for the transcriptome and proteome for noise level.

distributed with means varied based on noise level and a standard deviation of $\frac{1}{5}$ of the mean [3] [4]. These means vary between $\frac{1}{10}$ to $\frac{5}{10}$, with each level $\frac{1}{10}$ higher than the next, creating 5 total levels of noise. Secondly, after the noise amplitude percentage for each gene and sample is determined, we multiply this amplitude percentage by the assigned amplitude for the given gene. For linear and exponential models, this is 1 and its own amplitude, respectively. For oscillatory models, this is calculated by removing the baseline (constant $[y]$ for ECHO, linear $[\alpha t + y]$ for ECHO linear) and dividing the resulting maximum by 2.

In order to accurately test MOSAIC's joint modeling, we also needed to consider the different levels of noise between the transcriptome and proteome. To determine this exact separation, we observed the distributions of standard deviations of residuals of oscillatory genes found by MOSAIC in *Neurospora* [9, 10] (Figure 3). When we look at these distributions, we see that they are not completely overlapping, indicating that there should be a separation between RNA and protein (Figure 3A). Looking at the one-to-one difference between these standard deviations, we can also see that, while protein is higher, this difference is relatively normally distributed and centered on $\frac{2}{10}$ (Figure 3B). Thus, we adjusted the noise levels in proteome to be $\frac{2}{10}$ higher than the transcriptome, terming each separation a "set" and varying this from low to high (Figure 3C). This is detailed in Table 2.

[3] To prevent the generation of negative gene standard deviations, which are impossible, we restrict the range of this normal distribution to be in the range $\mu \pm 1$, where $\mu$ is the mean. Points outside of this range occur with almost 0 probability, so the normal distribution is not intensely disturbed.

[4] This $\frac{1}{5}$ ratio was chosen by computing the ratio of mean to standard deviations of the standard deviation of residuals in both the transcriptome and proteome in *Neurospora crassa* [9, 10], which both resulted in this ratio.

#### 4.2 Synthetic Data Analysis Methods

To analyze MOSAIC's accuracy fully, we took several approaches. First, to analyze its efficacy in model recovery, we we began by estimating AUCs for each model type using the ROC (Figure M.2A, Tables 3 to 8). Area Under the Curve (AUC) scores were calculated using the `roc.curve` function from the `PRROC` package [11, 8], for which we used 1 minus the BH-adjusted p-values. This indicated that a lower p-value had higher confidence in the model. If the model was misclassified, the BH-adjusted p-value was assigned to 1. We also looked at the percentages of each model that were misclassified and the missed models that made up this misclassification (Figures 4 to 6). In the resulting stacked bar plot, each of the true models appear on the x axis, while the resulting percentage of models misclassified or having a p-value higher than the cutoff make up the stacked y-axis.

To assess MOSAIC's joint modeling, we created a heat map of how many true oscillatory genes were recovered by joint modeling (Figure M.2B, 7). This difference was assessed by observing the selected models and BH-adjusted p-values with only model selection (before joint modeling) and after adding joint modeling (the full MOSAIC). Annotations within each box indicate the total before joint modeling, total after joint modeling, and the difference, separated by vertical bars.

Since we could reduce MOSAIC's results to a case of recovering oscillatory genes to nonoscillatory genes, we also decided to compare MOSAIC to other circadian-recovery methods: ECHO, JTK_CYCLE, and MetaCycle. All methods were run without preprocessing and with all indicated default settings. While typically an ROC curve would be used to determine success in this situation, we found that this balanced getting true positives and true negatives equally, which is not the case when evaluating circadian data. In the circadian situation, we care much more about the positives (obtaining a true circadian gene) than the negatives. Thus, instead we used F1-Scores and FDR/TPR comparisons.

We then use F1-Scores to estimate overall accuracy. The F1-score balances both the precision and recall of a test to compute a score in the following manner:

$$F_1 = 2\frac{\text{precision} \cdot \text{recall}}{\text{precision} + \text{recall}} \tag{13}$$

where precision and recall are defined as:

1. precision $= \frac{\text{TP}}{\text{TP+FP}}$

2. recall $= \frac{\text{TP}}{\text{FN+TP}}$

where T is true, F is false, P is positive, and N is negative. The highest The F1-score is ideal for the circadian problem, where we have an unbalanaced dataset with more noncircadian genes than circadian genes. We evaluated the F1-Score at BH-adjusted p-value cutoffs ranging from 0 to .1, encapsulating the range of common cutoffs. This formed a "curve" of accuracies, with the F1-Score's highest possible value at 1, and lowest at 0 (Figures M.3A, 8 to 10).

In FDR/TPR comparisons, we evaluated the false discovery rates (FDR) and true positive rates (TPR) of our methods at various common BH-adjusted p-value cutoffs (0.01, 0.05, 0.1) (Figures M.3B, 11 to 10). We then plotted (1-FDR) versus TPR; as you move along the diagonal of the plot towards the top right corner, the method becomes more accurate, balancing false discovery with true positives. This comparison shows each method's trade off of these quantities.

#### 4.3 Biological Analysis in *Neurospora*

In order to demonstrate MOSAIC's efficacy on real data, we applied our method to publicly available transcriptomic and proteomic data in *Neurospora crassa* [9, 10]. While other proteomic analyses exist [14, 16, 13], rhythmic protein detection in these studies were limited to often less than 1% of the proteome, which signals technical and analytical issues that preclude deep analysis. The Hurley et al. (2018) data, which features a time course over 48 hours with 2 hour resolution and 3 replicates and was analyzed with recent technological advances, lends itself more accurately to our analysis.

We used LIMBR to impute missing values and remove batch effects in the data as described in Hurley et al. (2018) [10, 3]. When running data through the MOSAIC interface, we used the preprocessing steps of normalization, removal of unexpressed genes, and smoothing (Section 3).

We also only looked for rhythms betweeen 20 to 24 hours, centered around the 22.5 hour rhythm for *Neurospora* [15].

To further understand this discrepancy between the role of circadian and non-circadian genes in the transcriptome and proteome of *Neurospora*, we also performed gene ontology (GO) analysis [2]. We followed the process proposed in ENCORE, an automatic circadian gene ontology enrichment application [5]. However, we changed our enrichment categories to be our significant circadian and non-circadian sets in the transcriptome and proteome, rather than the Amplitude Change coefficient categories built into the program. We used the Biological Process ontology and BH-adjusted p-values with a cutoff of 0.05 for enrichment, using all other program defaults. Enriched GO terms were categorized according to their closest second level parent, calculated using the `shortest_paths` function in `igraph` package [4]. For bar plots with third-level terms, a negative log BH-adjusted p-value indicates a BH-adjusted p-value of 1 for the corresponding term or that the term was pruned for nonsignificance by ENCORE. BH-adjusted p-values in the bar plot were also truncated if smaller than $1.5e-7$ ($-\log(p) \approx 15.7$), for ease of viewing. Exact enrichments for all GO terms appear in Supplemental File Tables 1 - 4.

##### 4.4 Software Version Notes

All specified data was run with the MOSAIC interface v0.2, the ECHO interface v3.21, JTK_CYCLE v3.1, MetaCycle v1.1.0, and ENCORE v3.0.3 unless otherwise noted. MOSAIC version updates and specifications can be found on GitHub [5].

#### 5 Synthetic Data Results Auxiliary Figures and Tables

##### 5.1 Model AUCs

AUCs for all MOSAIC models (Linear, Exponential, ECHO, ECHO Linear) appear in Tables 3 to 8. Further description of methods and data appears in Section 4.2.

Table 3: **AUCs for all MOSAIC models at varying conditions, fixed at one replicate in the transcriptome.** AUCs for all MOSAIC models at conditions varying noise and resolution for synthetic data in transcriptome and proteome, fixed at one replicate. Lin. = Linear, Exp. = Exponential, ECHO = ECHO, ECHO Lin. = ECHO Linear.

| | | Transcriptome, Replicates: 1 | | | | | | | | | | | |
|---|---|---|---|---|---|---|---|---|---|---|---|---|---|
| | | Resolution (Hours) | | | | | | | | | | | |
| | | 2 | | | | 4 | | | | 6 | | | |
| | | Lin. | Exp. | ECHO | ECHO Lin. | Lin. | Exp. | ECHO | ECHO Lin. | Lin. | Exp. | ECHO | ECHO Lin. |
| Noise | Low Set | 0.9478 | 0.96 | 0.9088 | 0.8746 | 0.8894 | 0.9188 | 0.8276 | 0.851 | 0.8199 | 0.8663 | 0.7275 | 0.8227 |
| | Medium Set | 0.9435 | 0.9196 | 0.9024 | 0.828 | 0.8728 | 0.8769 | 0.833 | 0.798 | 0.7622 | 0.808 | 0.7253 | 0.7561 |
| | High Set | 0.9282 | 0.8621 | 0.8989 | 0.7969 | 0.8561 | 0.8352 | 0.8172 | 0.7541 | 0.7369 | 0.7825 | 0.7084 | 0.7132 |

Table 4: **AUCs for all MOSAIC models at varying conditions, fixed at one replicate in the proteome.** AUCs for all MOSAIC models at conditions varying noise and resolution for synthetic data in proteome, fixed at one replicate. Lin. = Linear, Exp. = Exponential, ECHO = ECHO, ECHO Lin. = ECHO Linear.

| | | Proteome, Replicates: 1 | | | | | | | | | | | |
|---|---|---|---|---|---|---|---|---|---|---|---|---|---|
| | | Resolution (Hours) | | | | | | | | | | | |
| | | 2 | | | | 4 | | | | 6 | | | |
| | | Lin. | Exp. | ECHO | ECHO Lin. | Lin. | Exp. | ECHO | ECHO Lin. | Lin. | Exp. | ECHO | ECHO Lin. |
| Noise | Low Set | 0.9304 | 0.866 | 0.8953 | 0.7919 | 0.8379 | 0.8321 | 0.8274 | 0.7497 | 0.7311 | 0.7938 | 0.7195 | 0.7152 |
| | Medium Set | 0.9343 | 0.8338 | 0.8952 | 0.7616 | 0.8366 | 0.8047 | 0.8221 | 0.7123 | 0.7091 | 0.7825 | 0.7063 | 0.6719 |
| | High Set | 0.9326 | 0.8124 | 0.8773 | 0.7123 | 0.8255 | 0.7886 | 0.7971 | 0.684 | 0.6973 | 0.777 | 0.6903 | 0.6504 |

[5] https://github.com/delosh653/MOSAIC

Table 5: **AUCs for all MOSAIC models at varying conditions, fixed at two replicates in the transcriptome.** AUCs for all MOSAIC models at conditions varying noise and resolution for synthetic data in transcriptome, fixed at two replicates. Lin. = Linear, Exp. = Exponential, ECHO = ECHO, ECHO Lin. = ECHO Linear.

| | | **Transcriptome, Replicates: 2** | | | | | | | | | | | |
|---|---|---|---|---|---|---|---|---|---|---|---|---|---|
| | | **Resolution (Hours)** | | | | | | | | | | | |
| | | **2** | | | | **4** | | | | **6** | | | |
| | | **Lin.** | **Exp.** | **ECHO** | **ECHO Lin.** | **Lin.** | **Exp.** | **ECHO** | **ECHO Lin.** | **Lin.** | **Exp.** | **ECHO** | **ECHO Lin.** |
| **Noise** | **Low Set** | 0.9819 | 0.9441 | 0.866 | 0.8776 | 0.9893 | 0.9274 | 0.9148 | 0.8145 | 0.9849 | 0.93 | 0.8813 | 0.7743 |
| | **Medium Set** | 0.9778 | 0.8533 | 0.8597 | 0.8301 | 0.9815 | 0.8332 | 0.8868 | 0.799 | 0.9708 | 0.8443 | 0.8574 | 0.7681 |
| | **High Set** | 0.9764 | 0.7978 | 0.8527 | 0.8048 | 0.9838 | 0.7927 | 0.8669 | 0.769 | 0.9582 | 0.7957 | 0.8264 | 0.7474 |

Table 6: **AUCs for all MOSAIC models at varying conditions, fixed at two replicates in the proteome.** AUCs for all MOSAIC models at conditions varying noise and resolution for synthetic data in proteome, fixed at two replicates. Lin. = Linear, Exp. = Exponential, ECHO = ECHO, ECHO Lin. = ECHO Linear.

| | | **Proteome, Replicates: 2** | | | | | | | | | | | |
|---|---|---|---|---|---|---|---|---|---|---|---|---|---|
| | | **Resolution (Hours)** | | | | | | | | | | | |
| | | **2** | | | | **4** | | | | **6** | | | |
| | | **Lin.** | **Exp.** | **ECHO** | **ECHO Lin.** | **Lin.** | **Exp.** | **ECHO** | **ECHO Lin.** | **Lin.** | **Exp.** | **ECHO** | **ECHO Lin.** |
| **Noise** | **Low Set** | 0.9809 | 0.8019 | 0.8473 | 0.8009 | 0.9828 | 0.7918 | 0.8654 | 0.7672 | 0.9578 | 0.7997 | 0.8355 | 0.7398 |
| | **Medium Set** | 0.9733 | 0.7811 | 0.8432 | 0.7724 | 0.9784 | 0.7785 | 0.8467 | 0.736 | 0.9534 | 0.7827 | 0.8033 | 0.7111 |
| | **High Set** | 0.973 | 0.7685 | 0.827 | 0.7286 | 0.9767 | 0.7688 | 0.8262 | 0.6963 | 0.9558 | 0.7704 | 0.795 | 0.6901 |

Table 7: **AUCs for all MOSAIC models at varying conditions, fixed at three replicates in the transcriptome.** AUCs for all MOSAIC models at conditions varying noise and resolution for synthetic data in transcriptome, fixed at three replicates. Lin. = Linear, Exp. = Exponential, ECHO = ECHO, ECHO Lin. = ECHO Linear.

| | | **Transcriptome, Replicates: 3** | | | | | | | | | | | |
|---|---|---|---|---|---|---|---|---|---|---|---|---|---|
| | | **Resolution (Hours)** | | | | | | | | | | | |
| | | **2** | | | | **4** | | | | **6** | | | |
| | | **Lin.** | **Exp.** | **ECHO** | **ECHO Lin.** | **Lin.** | **Exp.** | **ECHO** | **ECHO Lin.** | **Lin.** | **Exp.** | **ECHO** | **ECHO Lin.** |
| **Noise** | **Low Set** | 0.9909 | 0.9876 | 0.9074 | 0.892 | 0.992 | 0.9859 | 0.9406 | 0.831 | 0.995 | 0.972 | 0.9064 | 0.7363 |
| | **Medium Set** | 0.9884 | 0.9423 | 0.9046 | 0.8526 | 0.9832 | 0.922 | 0.9246 | 0.8086 | 0.9881 | 0.881 | 0.8941 | 0.733 |
| | **High Set** | 0.9898 | 0.8727 | 0.9079 | 0.8302 | 0.9835 | 0.85 | 0.8983 | 0.7883 | 0.9785 | 0.8162 | 0.8808 | 0.7151 |

Table 8: **AUCs for all MOSAIC models at varying conditions, fixed at three replicates in the proteome.** AUCs for all MOSAIC models at conditions varying noise and resolution for synthetic data in proteome, fixed at three replicates. Lin. = Linear, Exp. = Exponential, ECHO = ECHO, ECHO Lin. = ECHO Linear.

| | | **Proteome, Replicates: 3** | | | | | | | | | | | |
|---|---|---|---|---|---|---|---|---|---|---|---|---|---|
| | | **Resolution (Hours)** | | | | | | | | | | | |
| | | **2** | | | | **4** | | | | **6** | | | |
| | | **Lin.** | **Exp.** | **ECHO** | **ECHO Lin.** | **Lin.** | **Exp.** | **ECHO** | **ECHO Lin.** | **Lin.** | **Exp.** | **ECHO** | **ECHO Lin.** |
| **Noise** | **Low Set** | 0.9881 | 0.8804 | 0.9024 | 0.8217 | 0.9783 | 0.8522 | 0.9014 | 0.7749 | 0.9745 | 0.8257 | 0.8791 | 0.7063 |
| | **Medium Set** | 0.9861 | 0.8297 | 0.8952 | 0.805 | 0.9814 | 0.8164 | 0.8969 | 0.7617 | 0.9689 | 0.7955 | 0.863 | 0.6953 |
| | **High Set** | 0.9875 | 0.8014 | 0.8903 | 0.7722 | 0.981 | 0.7923 | 0.8836 | 0.7287 | 0.9659 | 0.7804 | 0.8455 | 0.6644 |

#### 5.2 Misclassified Model Bar Plots

Bar plots of misclassified models for all MOSAIC models (Linear, Exponential, ECHO, ECHO Linear) appear in Figures 4 to 6. Further description of methods and data appears in Section 4.2.

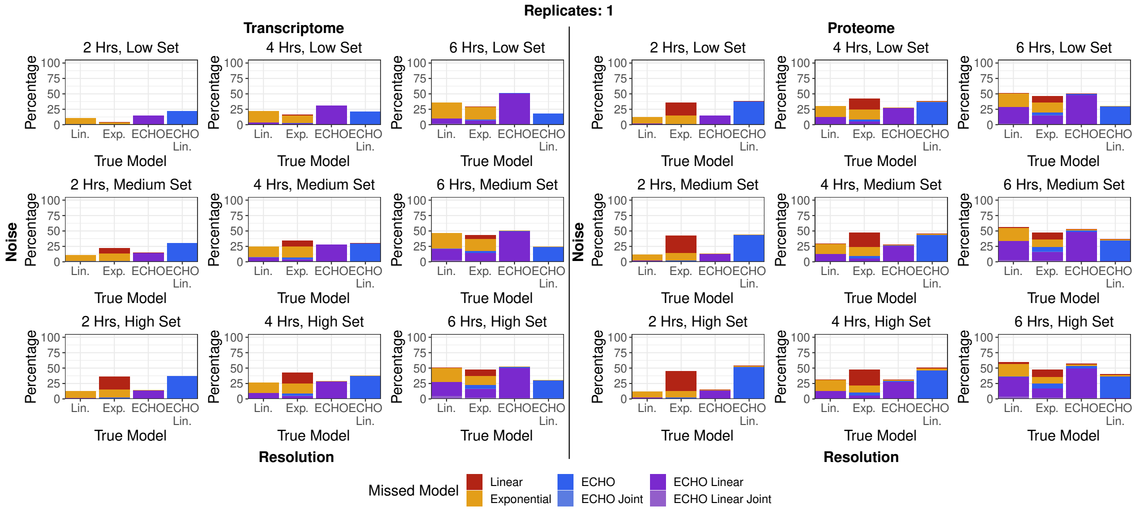


Figure 4: **Bar plots of misclassified models for all MOSAIC models, fixed at one replicate in both omics types.** Bar plot of the percentage of missed models for each model type, and what models these were misclassified to. Conditions were varied in noise and resolution for synthetic data in transcriptome and proteome, fixed at one replicate. Hrs = Hours. Lin. = Linear, Exp. = Exponential, ECHO = ECHO, ECHO Lin. = ECHO Linear.

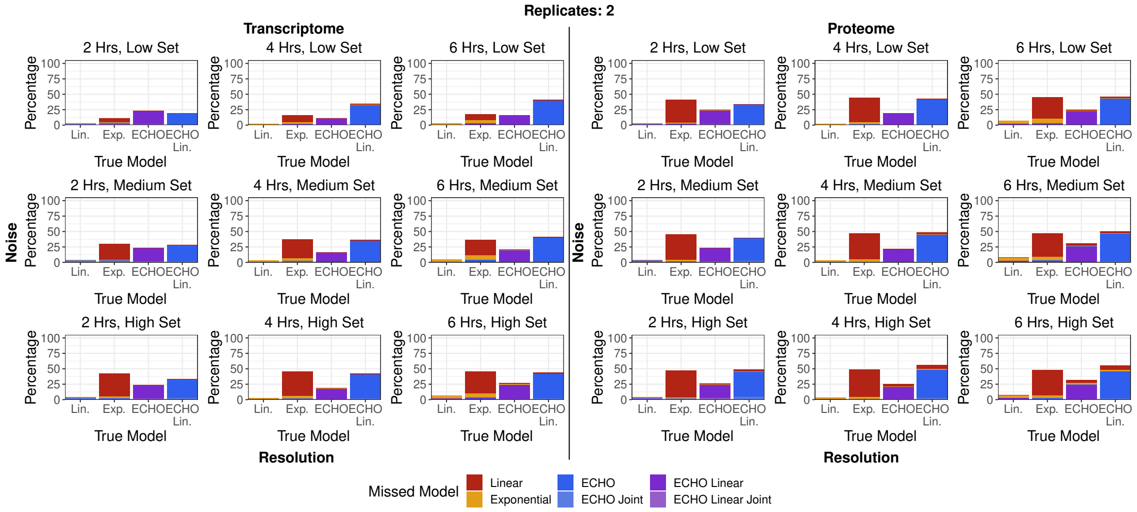


Figure 5: **Bar plots of misclassified models for all MOSAIC models, fixed at two replicates in both omics types.** Bar plot of the percentage of missed models for each model type, and what models these were misclassified to. Conditions were varied in noise and resolution for synthetic data in transcriptome and proteome, fixed at two replicates. Hrs = Hours. Lin. = Linear, Exp. = Exponential, ECHO = ECHO, ECHO Lin. = ECHO Linear.

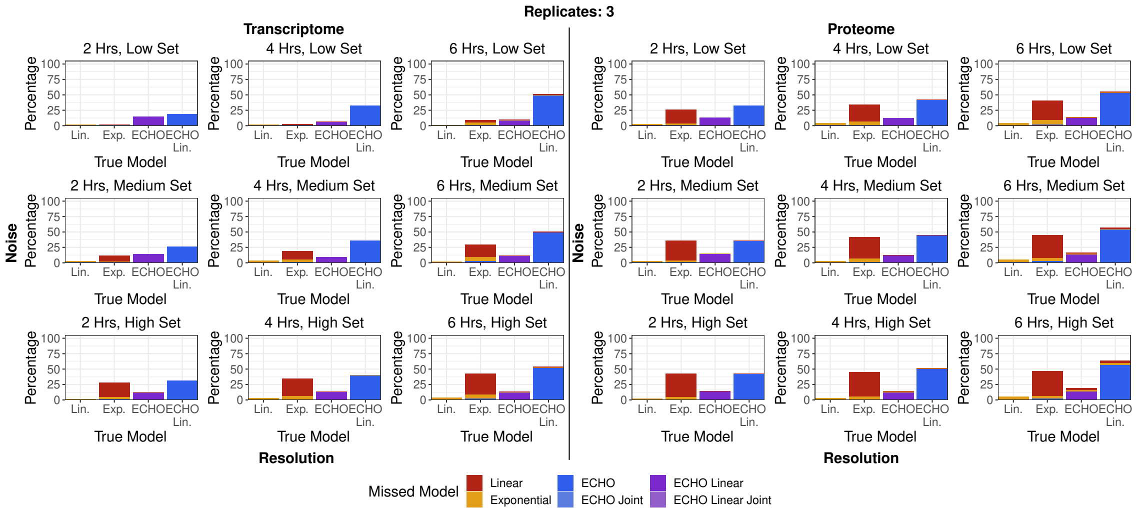


Figure 6: **Bar plots of misclassified models for all MOSAIC models, fixed at three replicates in both omics types.** Bar plot of the percentage of missed models for each model type, and what models these were misclassified to. Conditions were varied in noise and resolution for synthetic data in transcriptome and proteome, fixed at three replicates. Hrs = Hours. Lin. = Linear, Exp. = Exponential, ECHO = ECHO, ECHO Lin. = ECHO Linear.

#### 5.3 Joint Modeling Oscillation Recovery

Heat maps of recovered true oscillatory genes after the addition of joint modeling in MOSAIC, at varying conditions in in synthetic transcriptome and proteome synthetic data appear in Figure 7. Further description of methods and data appears in Section 4.2.

#### 5.4 F1-Score Curves

F1-score curves, computing F1-score at various BH-adjusted p-value cutoffs, for all tested methods (ECHO, JTK_CYCLE, MetaCycle, MOSAIC) appear in Figs. 8 to 10. Further description of methods and data appears in Section 4.2.

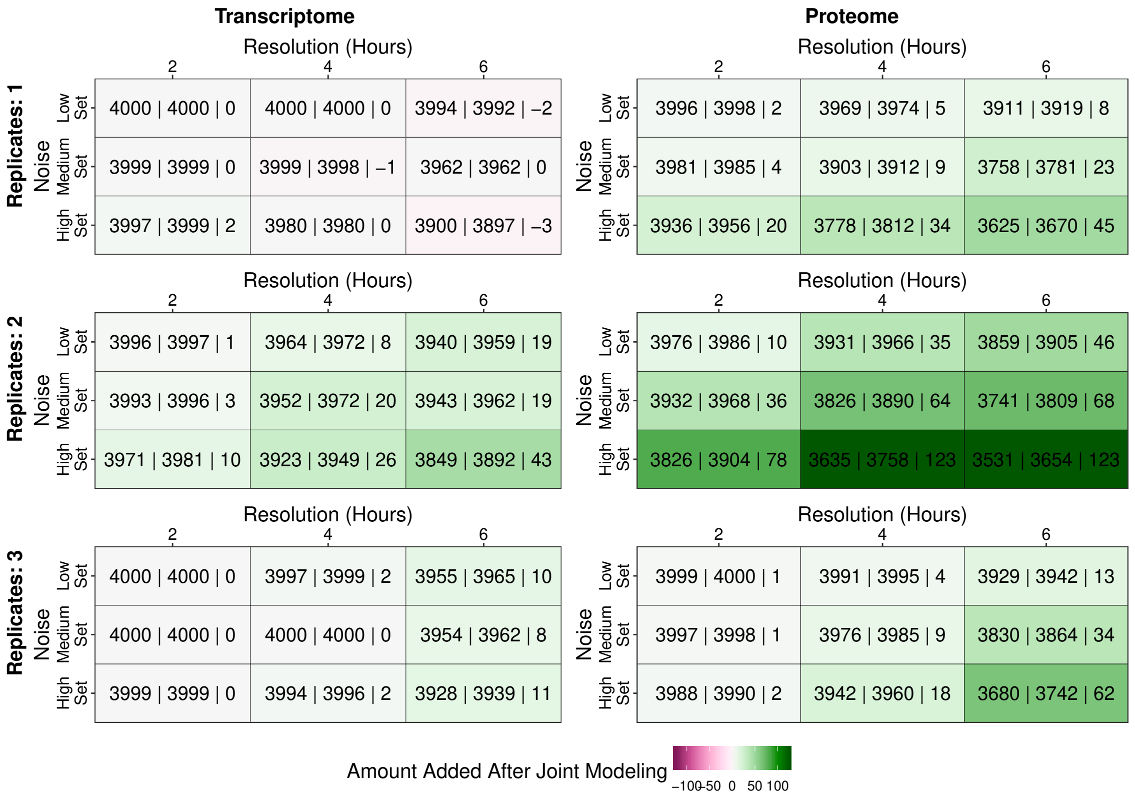


Figure 7: **MOSAIC's joint modeling recovers large amounts of data in both the transcriptome and proteome.** Heat maps of recovered true oscillatory genes after the addition of joint modeling in MOSAIC, at varying conditions in synthetic transcriptome and proteome synthetic data. Annotations in each condition indicate the amount of true oscillatory genes recovered before joint modeling, after joint modeling, and the difference, with vertical lines separating each amount. Methods described in Section 4.2.

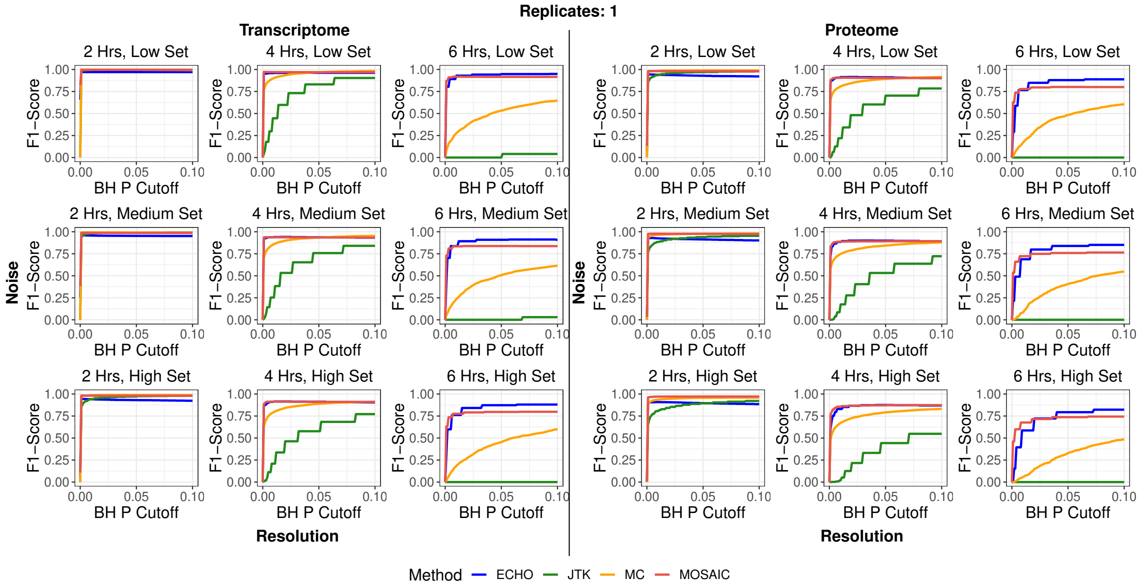


Figure 8: **F1-score curves for various methods at varying conditions, fixed for one replicate for both omics types.** F1-score curves evaluating F1-score at cutoffs varying between 0 and 0.1 for all tested methods. Conditions were varied in noise and resolution for synthetic data in transcriptome and proteome, fixed at one replicate. Hrs = Hours. ECHO = ECHO, JTK = JTK_CYCLE, MC = MetaCycle, MOSAIC = MOSAIC.

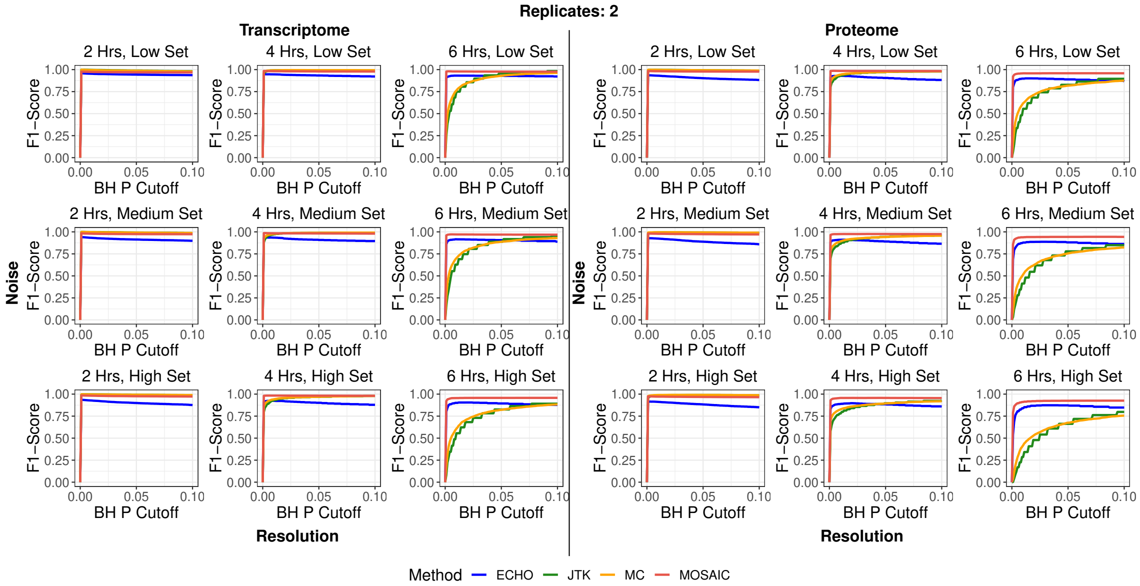


Figure 9: **F1-score curves for various methods at varying conditions, fixed for two replicates for both omics types.** F1-score curves evaluating F1-score at cutoffs varying between 0 and 0.1 for all tested methods. Conditions were varied in noise and resolution for synthetic data in transcriptome and proteome, fixed at two replicates. Hrs = Hours. ECHO = ECHO, JTK = JTK_CYCLE, MC = MetaCycle, MOSAIC = MOSAIC.

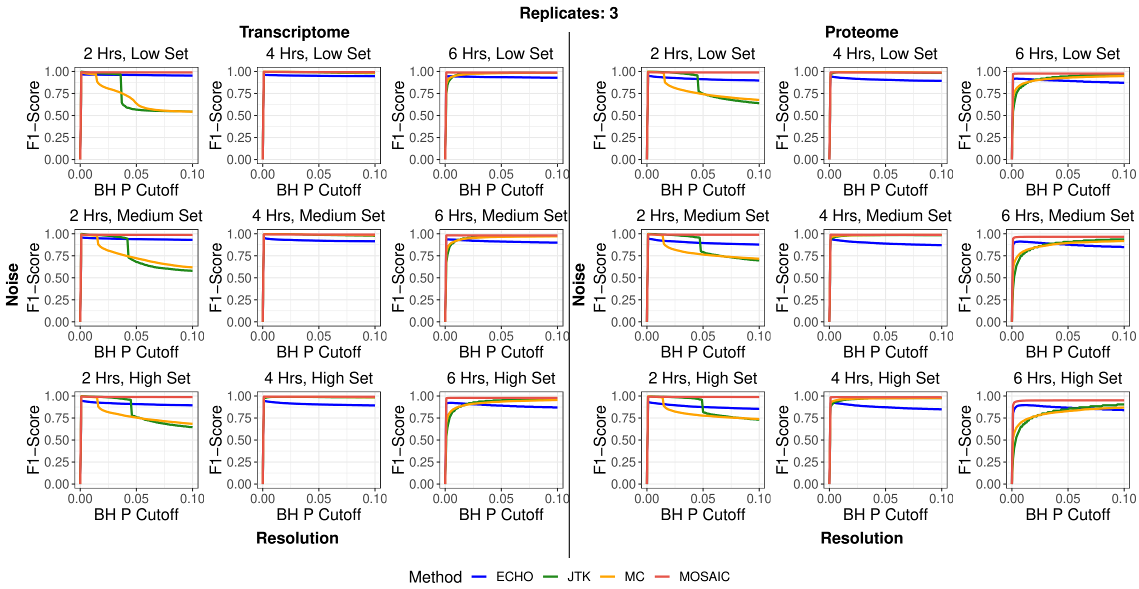

Figure 10: **F1-score curves for various methods at varying conditions, fixed for three replicates for both omics types.** F1-score curves evaluating F1-score at cutoffs varying between 0 and 0.1 for all tested methods. Conditions were varied in noise and resolution for synthetic data in transcriptome and proteome, fixed at three replicates. Hrs = Hours. ECHO = ECHO, JTK = JTK_CYCLE, MC = MetaCycle, MOSAIC = MOSAIC.

#### 5.5 TPR/FDR Comparisons

Comparison scatter plots of FDR versus TPR, computed at various cutoffs for all tested methods, appear in Figs. 11 to 13. Further description of methods and data appears in Section 4.2.

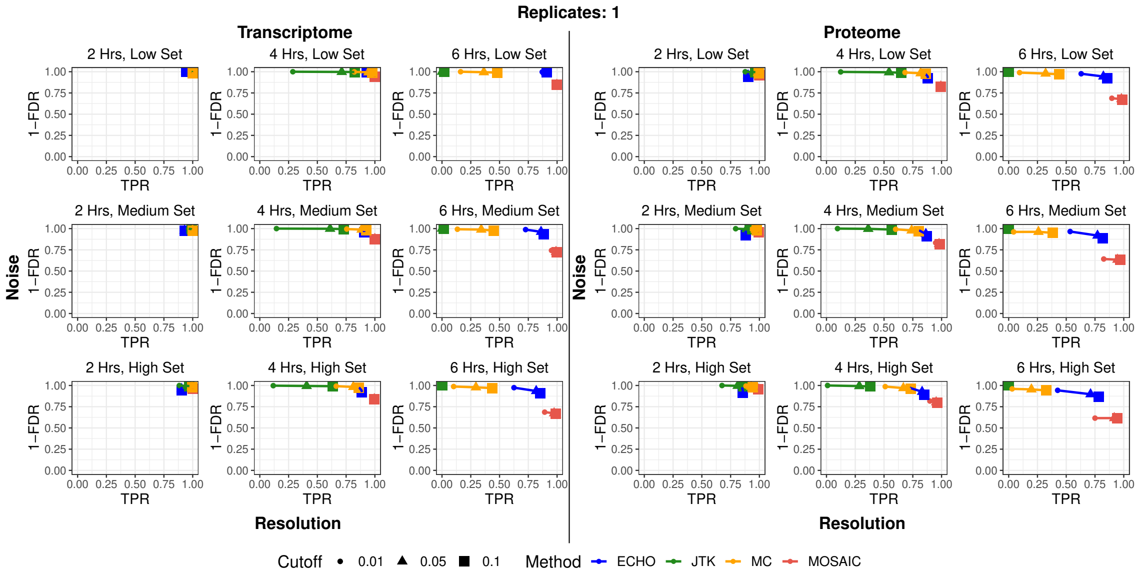

Figure 11: **Comparisons between TPR and FDR for all tested methods at various conditions, fixed at one replicate for both omics types.** Scatter plots of (1-FDR) versus FDR at varying BH-adjusted p-value cutoffs (0.01, 0.05 0.1) for all tested methods. Conditions were varied in noise and resolution for synthetic data in transcriptome and proteome, fixed at one replicate. Hrs = Hours. ECHO = ECHO, JTK = JTK_CYCLE, MC = MetaCycle, MOSAIC = MOSAIC.

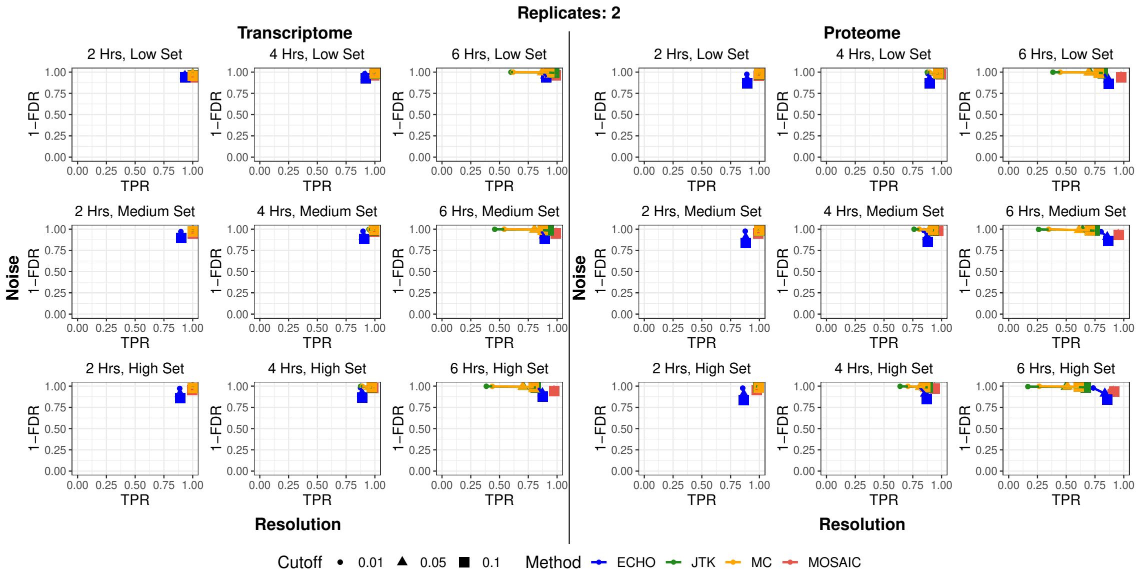


Figure 12: **Comparisons between TPR and FDR for all tested methods at various conditions, fixed at two replicates for both omics types.** Scatter plots of (1-FDR) versus FDR at varying BH-adjusted p-value cutoffs (0.01, 0.05 0.1) for all tested methods. Conditions were varied in noise and resolution for synthetic data in transcriptome and proteome, fixed at two replicates. Hrs = Hours. ECHO = ECHO, JTK = JTK_CYCLE, MC = MetaCycle, MOSAIC = MOSAIC.

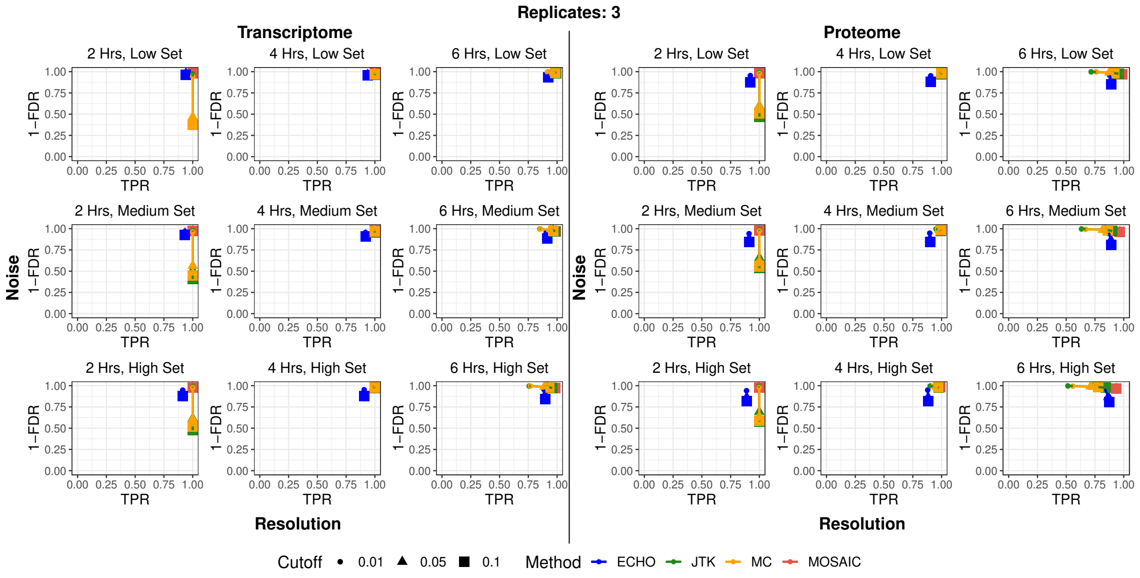


Figure 13: **Comparisons between TPR and FDR for all tested methods at various conditions, fixed at three replicates for both omics types.**. Scatter plots of (1-FDR) versus FDR at varying BH-adjusted p-value cutoffs (0.01, 0.05 0.1) for all tested methods. Conditions were varied in noise and resolution for synthetic data in transcriptome and proteome, fixed at three replicates. Hrs = Hours. ECHO = ECHO, JTK = JTK_CYCLE, MC = MetaCycle, MOSAIC = MOSAIC.

### References

[1] H. Akaike. A new look at the statistical model identification. *IEEE Trans. Autom. Control*, 19(6):716–723, December 1974.

[2] Michael Ashburner, Catherine A. Ball, Judith A. Blake, David Botstein, Heather Butler, J. Michael Cherry, Allan P. Davis, Kara Dolinski, Selina S. Dwight, Janan T. Eppig, Midori A. Harris, David P. Hill, Laurie Issel-Tarver, Andrew Kasarskis, Suzanna Lewis, John C. Matese, Joel E. Richardson, Martin Ringwald, Gerald M. Rubin, and Gavin Sherlock. Gene ontology: tool for the unification of biology. *Nat. Genet.*, 25(1):25–29, May 2000.

[3] Alexander M Crowell, Casey S Greene, Jennifer J Loros, and Jay C Dunlap. Learning and imputation for mass-spec bias reduction (LIMBR). *Bioinf.*, 35(9):1518–1526, September 2018.

[4] Gabor Csardi and Tamas Nepusz. The igraph software package for complex network research. *InterJournal*, Complex Systems:1695, 2006.

[5] Hannah De los Santos, Kristin P. Bennett, and Jennifer M. Hurley. ENCORE: A visualization tool for insight into circadian omics. In *Proc. of the 10th ACM Int. Conf. on Bioinf., Comput. Biol. and Health Inform.*, ACM-BCB '19, New York, NY, USA, 2019. Niagara Falls, NY, USA, ACM.

[6] Hannah De los Santos, Emily J Collins, Catherine Mann, April W Sagan, Meaghan S Jankowski, Kristin P Bennett, and Jennifer M Hurley. ECHO: an application for detection and analysis of oscillators identifies metabolic regulation on genome-wide circadian output. *Bioinf.*, 36(3):773–781, February 2020.

[7] Anastasia Deckard, John Harer, John B. Hogenesch, Ron C. Anafi, and Steven B. Haase. Design and analysis of large-scale biological rhythm studies: a comparison of algorithms for detecting periodic signals in biological data. *Bioinf.*, 29(24):3174–3180, September 2013.

[8] Jan Grau, Ivo Grosse, and Jens Keilwagen. Prroc: computing and visualizing precision-recall and receiver operating characteristic curves in r. *Bioinf.*, 31(15):2595–2597, 2015.

[9] Jennifer M. Hurley, Arko Dasgupta, Jillian M. Emerson, Xiaoying Zhou, Carol S. Ringelberg, Nicole Knabe, Anna M. Lipzen, Erika A. Lindquist, Christopher G. Daum, Kerrie W. Barry, Igor V. Grigoriev, Kristina M. Smith, James E. Galagan, Deborah Bell-Pedersen, Michael Freitag, Chao Cheng, Jennifer J. Loros, and Jay C. Dunlap. Analysis of clock-regulated genes in Neurospora reveals widespread posttranscriptional control of metabolic potential. *Proc. Natl. Acad. Sci.*, 111(48):16995–17002, October 2014.

[10] Jennifer M. Hurley, Meaghan S. Jankowski, Hannah De los Santos, Alexander M. Crowell, Samuel B. Fordyce, Jeremy D. Zucker, Neeraj Kumar, Samuel O. Purvine, Errol W. Robinson, Anil Shukla, Erika Zink, William R. Cannon, Scott E. Baker, Jennifer J. Loros, and Jay C. Dunlap. Circadian proteomic analysis uncovers mechanisms of post-transcriptional regulation in metabolic pathways. *Cell Syst.*, 7(6):613–626.e5, December 2018.

[11] Jens Keilwagen, Ivo Grosse, and Jan Grau. Area under precision-recall curves for weighted and unweighted data. *PLoS ONE*, 9(3):e92209, March 2014.

[12] Jonathan M. Raser and Erin K. O'Shea. Noise in gene expression: Origins, consequences, and control. *Science*, 309(5743):2010–2013, 2005.

[13] Akhilesh B. Reddy, Natasha A. Karp, Elizabeth S. Maywood, Elizabeth A. Sage, Michael Deery, John S. O'Neill, Gabriel K.Y. Wong, Jo Chesham, Mark Odell, Kathryn S. Lilley, Charalambos P. Kyriacou, and Michael H. Hastings. Circadian orchestration of the hepatic proteome. *Curr. Biol.*, 16(11):1107–1115, June 2006.

[14] Maria S. Robles, Jürgen Cox, and Matthias Mann. In-vivo quantitative proteomics reveals a key contribution of post-transcriptional mechanisms to the circadian regulation of liver metabolism. *PLoS Genet.*, 10(1):e1004047, January 2014.

[15] M. L. Sargent, W. R. Briggs, and D. O. Woodward. Circadian nature of a rhythm expressed by an invertaseless strain of neurospora crassa. *Plant Physiol.*, 41(8):1343–1349, October 1966.

[16] Jingkui Wang, Daniel Mauvoisin, Eva Martin, Florian Atger, Antonio Núñez Galindo, Loïc Dayon, Federico Sizzano, Alessio Palini, Martin Kussmann, Patrice Waridel, Manfredo Quadroni, Vjekoslav Dulić, Felix Naef, and Frédéric Gachon. Nuclear proteomics uncovers diurnal regulatory landscapes in mouse liver. *Cell Metab.*, 25(1):102–117, January 2017.

[17] Pål O. Westermark, David K. Welsh, Hitoshi Okamura, and Hanspeter Herzel. Quantification of Circadian Rhythms in Single Cells. *PloS Comput. Biol*, 5(11):e1000580, November 2009.

[18] Gang Wu, Ron C. Anafi, Michael E. Hughes, Karl Kornacker, and John B. Hogenesch. MetaCycle: an integrated r package to evaluate periodicity in large scale data. *Bioinf.*, 32(21):3351–3353, July 2016.
